# Quantitative analysis of synaptic zinc in the brain by covalent chemistry-based semisynthetic biosensors

**DOI:** 10.64898/2025.12.22.695889

**Authors:** Hao Zhu, Seiji Sakamoto, Keisuke Nakamura, Qiu Chengli, Kohei Nakajima, Hiroshi Nonaka, Itaru Hamachi

## Abstract

Labile zinc ions (Zn²⁺) are stored in glutamatergic vesicles at specific excitatory synapses in the central nervous system and released into synaptic clefts in an activity-dependent manner. Although Zn²⁺ is suggested to modulate various neuroreceptor functions, its precise roles remain unclear due to a lack of tools capable of quantitatively analyzing Zn²⁺ with synapse-level spatial resolution. Here, we developed neuroreceptor-based semisynthetic sensors that record synaptic Zn²⁺ dynamics by covalent chemistry in the living mouse brain. Using a chemical knock-in strategy, we introduced activity-based Zn^2+^-probes with distinct affinities into endogenous α-amino-3-hydroxy-5-methyl-4-isoxazole-propionic acid or γ-aminobutyric acid type A receptors. These in-brain-constructed Zn²⁺ sensors enabled, to our knowledge, for the first time, quantitative, region-specific mapping of Zn²⁺ released into synaptic clefts. Imaging-based analyses revealed differences in Zn²⁺ concentrations at excitatory and inhibitory synapses across hippocampal regions during kainate-induced seizures, providing new insights into the physiological and pathological functions of synaptic Zn²⁺.

## Introduction

Zinc is an essential trace element that plays a variety of structural, catalytic and signaling roles in biological systems. In the brain, a specialized pool of free or loosely bound labile zinc ions (Zn^2+^) is sequestered in glutamatergic vesicles via the vesicular Zn^2+^ transporter (ZnT3) at many excitatory synapses, particularly in areas such as the hippocampus, cerebral cortex and amygdala.^1–4^ Upon neuronal excitation, Zn^2+^ is co-released with glutamate into the synaptic cleft, where it is thought to interact with and allosterically modulate various neurotransmitter receptors (neuroreceptors), including N-methyl-D-aspartate receptors (NMDARs),^5^ α-amino-3-hydroxy-5-methyl-4-isoxazole-propionic acid receptors (AMPARs),^6^ and γ-aminobutyric acid receptors (GABA_A_Rs).^7^ Synaptically released Zn^2+^ is considered to regulate both excitatory and inhibitory synaptic transmission and contribute to synaptic plasticity. However, dysregulated Zn^2+^ signaling has been implicated in excitotoxic conditions, such as epilepsy, ischemia and traumatic brain injury, as well as in the pathogenesis of neurodegenerative disorders, including Alzheimer’s disease.^4^ Despite such importance, the precise roles of Zn^2+^ in the central nervous system have not been systematically elucidated at the molecular level, because of the lack of analytical tools applicable to the living brain.

In addition to traditional histochemical methods (e.g., Timm staining^8^) and microdialysis,^9^ a few invaluable biochemical tools including Zn^2+^ chelators,^10^ ZnT3 knockout mice,^11^ fluorescent Zn^2+^ sensors^12–14^ have been used for studying the biological roles of Zn^2+^ in the brain to date, and sought to map the spatial distribution of synaptic Zn^2+^ and monitor its dynamics in the nervous system. Lippard and colleagues developed an elegant method that involves loading a membrane-impermeant Zn^2+^ sensor into presynaptic vesicles via KCl-induced depolarization, allowing for visualization of Zn^2+^ release at individual presynaptic sites.^15^ Although powerful, application of these small-molecule sensors has been largely limited to cultured neurons and acutely prepared brain slice experiments,^16^ rather than to the living brain. Moreover, it was previously pointed out that up to 50% of the synaptic Zn^2+^ is lost during the conventional hippocampal slice preparation.^17^ Genetically encoded membrane-bound Zn^2+^ sensors were developed for imaging activity-dependent extracellular Zn^2+^ release in the living mouse brain.^18^ However, these techniques suffer from insufficient spatial resolution, difficulty in addressing deep-tissue imaging and poor quantitative determination of Zn^2+^ concentrations. There is now a strong demand for methods that can analyze transient changes in Zn^2+^ dynamics at high spatial resolution within the structurally delicate 3D-environment of the brain.

Converting such transient biological events into permanent detection signals aids their precise analysis. Activity-based labeling is a valuable strategy used to transform transient analyte activities into chemical bond formation (or cleavage), thereby enabling later in-depth analyses such as high-resolution imaging and immunohistochemistry. The utility of such labeling has been demonstrated in studies of intracellular dynamics of reactive oxygen species, calcium and transient metal ions such as zinc and copper.^19–26^ A recording strategy based on covalent chemistry for capturing transient events could serve as an even more powerful methodology for analysis in vivo, including those performed in the brain, where structural complexity poses substantial challenges. Johnsson and co-workers recently reported split-Halo Tag systems whose self-labeling with chloroalkane-fluorophore substrates occurs only in the presence of specific, transient cellular events.^27,28^ These genetically engineered Halo Tag-based sensors enabled the recording of various transient events such as protein-protein interactions, G-protein-coupled receptor activation, increases in intracellular Ca^2+^ levels and kinase activities, by permanent covalent bond formation not only in living cells but also in living animals including fruit flies (*Drosophila*), zebrafishes and freely moving mice. These recordings provided a critical clue for covalent-chemistry-based Zn^2+^ sensors applicable to the living brain.

Here, we describe the development of neuroreceptor-based semisynthetic sensors that enable the recording of the synaptic Zn^2+^ dynamics through covalent bond cleavages and perform the quantitative analyses of Zn^2+^ in the living mouse brain (Fig. 1a). We designed a set of Zn^2+^-responsive probes with distinct affinities whose linker is irreversibly cleaved only in the presence of Zn^2+^, allowing us to record Zn^2+^ dynamics with a permanent signal. These probes were tethered to endogenous neuroreceptors located in either excitatory or inhibitory synapses using our recently established “chemical knock-in (KI)” protocol.^29,30^ Using these in-brain-constructed Zn^2+^ sensors, we achieved, for the first time to our knowledge, the quantitative and region-distinct evaluation of Zn^2+^ released into the synaptic clefts in the living mouse brain. A subsequent in-depth imaging-based quantitative analysis revealed that Zn^2+^ concentrations at excitatory synapses differ across hippocampal areas during kainate-induced seizures, and that transient Zn^2+^ concentrations differ between excitatory and inhibitory synapses even within the same brain region.

**Fig. 1.**
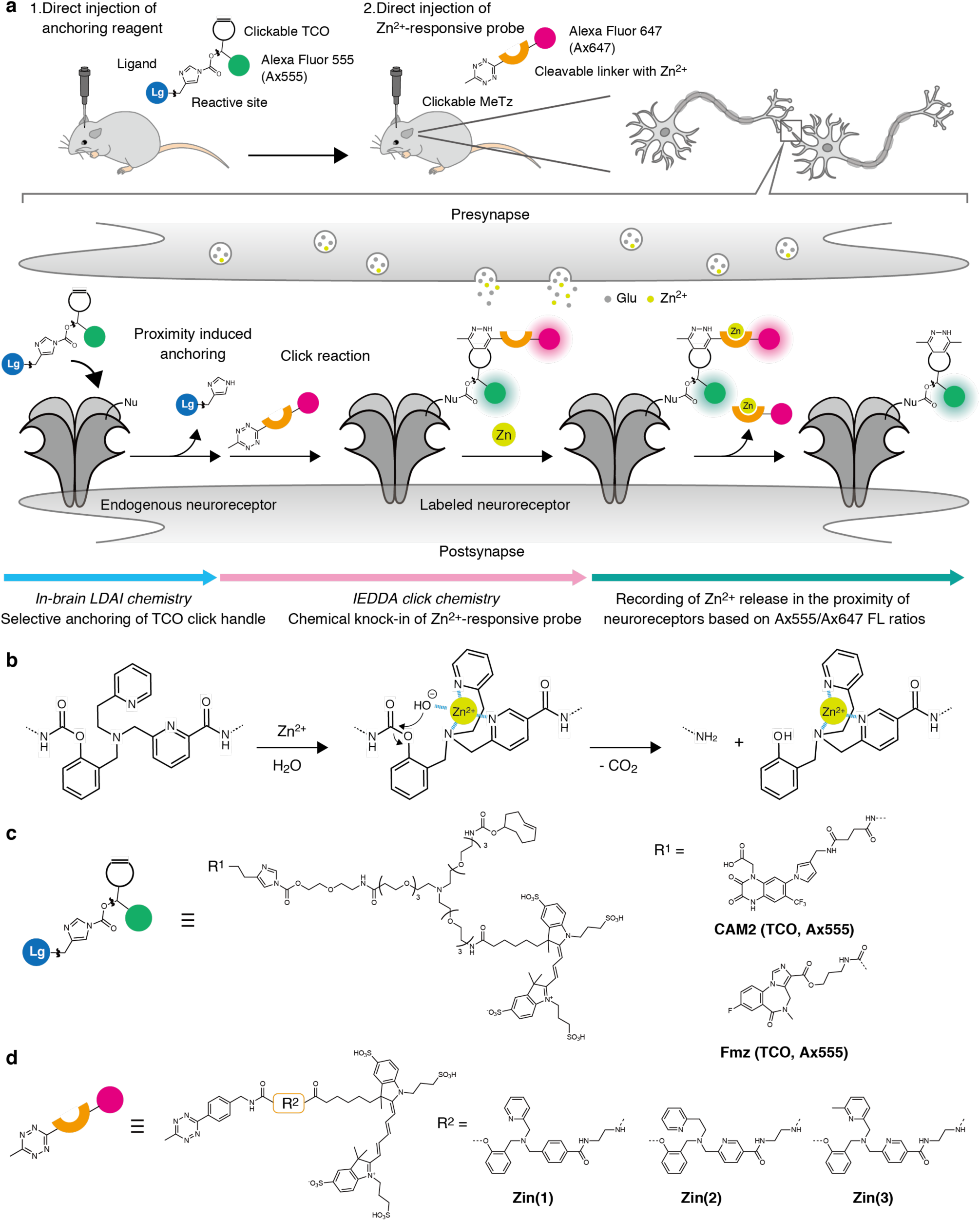
Chemical knock-in of Zn^2+^-responsive probes to neuroreceptors in a living mouse brain. **a**, Schematic illustration of a neuroreceptor-based fluorescent sensor and the recording of Zn^2+^ release at synapses. Nu, nucleophilic amino acid residue; Lg, selective ligand for target receptor; TCO, *trans*-cyclooctene; MeTz, methyltetrazine. **b**, A proposed mechanism of Zn^2+^-promoted hydrolysis of the carbamate linkage. **c**, Chemical structures of LDAI reagents for anchoring TCO to AMPARs and GABA_A_Rs. **d**, Chemical structures of Zn^2+^-responsive probes tethering MeTz group. The structures of Alexa Fluor 647 (Ax647) and Alexa Fluor 555 (Ax555) are putative structure.

## Results

### Characterization of Zn^2+^-promoted hydrolysis of the carbamate linkage in vitro

To record the transient Zn^2+^ signals in synapses in the living mouse brain, we tethered Zn^2+^-activity-based reaction probes to neuroreceptors located at specific synapses, which can generate semisynthetic sensors (Fig. 1a). Dipicolylamine derivatives were employed as Zn^2+^-chelating units, and a carbamate bond was selected as a cleavable linker that is hydrolytically degraded upon Zn^2+^-binding at the chelator site (Fig. 1b).^31,32^ Given that the bond cleavage is irreversible, our Zn^2+^ sensor can record the synaptic Zn^2+^ release using an integration detection mode. We chose two distinct receptors as scaffolds for semisynthetic fluorescent sensors of Zn^2+^, namely, AMPARs and GABA_A_Rs, which are endogenously expressed in excitatory and inhibitory synapses, respectively. A Zn^2+^-responsive module comprising a Zn^2+^-chelator and a fluorophore (Alexa Fluor 647 (Ax647)) linked through the carbamate bond is tethered to a target neuroreceptor using our chemical KI method.^29,30^ The chemical KI was conducted using ligand-directed acyl imidazole (LDAI) chemistry to modify a TCO handle and another fluorophore (Alexa Fluor 555 (Ax555)) as an internal standard, followed by an IEDDA click reaction with a MeTz-attached Zn^2+^ responsive module, by which visualization and quantitative analyses of the synaptically released Zn^2+^ proximal to the target receptor are effected (Fig. 1a). A previously reported anchoring reagent, **CAM2 (TCO, Ax555)** (referred to as **2 (Ax555)**^30^), was employed to incorporate the TCO handle and Ax555 fluorophore into the AMPARs. (Fig. 1c). As an anchoring reagent for GABA_A_Rs, **Fmz (TCO, Ax555)**, was newly designed by replacing the PFQX of **CAM2 (TCO, Ax555)** with flumazenil, a specific ligand for GABA_A_Rs with nM affinity (Fig. 1c). As Zn^2+^-responsive modules, we prepared a series of **Zinc probes** exhibiting different Zn^2+^ affinities, **Zin(1)**, **Zin(2)** and **Zin(3)**, where Ax647 and MeTz were connected via Zn^2+^-cleavable linkers (Fig. 1d).^16^

Before construction of neuroreceptors-based Zn^2+^ sensors, we firstly characterized Zn^2+^-promoted cleavage of the carbamate linkage in compounds **1, 2** and **3** as models of **Zinc probes Zin(1), Zin(2) and Zin(3)**, respectively, by high performance liquid chromatography (HPLC) (Fig. 2a). The chelated Zn^2+^ activates the coordination water due to its Lewis acidity, thereby the neighboring carbamate bond is readily hydrolyzed.^31,32^ Incubating **1, 2** or **3** with excess Zn^2+^ at 37 °C results in their efficient conversion to the corresponding hydrolyzed products (Fig. 2b, Supplementary Fig. 1 and Supplementary Fig. 2). By varying the Zn^2+^ concentrations, the EC_50_ values of Zn^2+^ for **1, 2** and **3** were determined as 7.7 × 10^−5^ M, 7.0 × 10^−7^ M and 2.4 ×10^−8^ M, respectively (Fig. 2c), which are close to the *K*_d_ values of fluorescent sensors possessing similar Zn^2+^ chelators.^16^ Thus, a combination of these sensor modules is expected to enable the detection of Zn^2+^ in a wide range of concentrations (10^−8^ to 10^−4^ M), which aligns with the previously estimated synaptic Zn^2+^ levels.^16,33–36^ Although 1 µM of Zn^2+^ dramatically promoted the hydrolysis of **2**, other biologically relevant metal ions, such as Ca^2+^ and Mg^2+^, were not effective, even at mM concentrations (Fig. 2d). Other transition metal ions, such as Co^2+^ and Cu^2+^, can also facilitate the hydrolysis of **2**, consistent with their reported hydrolytic abilities in small-molecule biomimetic systems.^37^ However, unlike Zn^2+^, these transition metals are less abundant in appreciable chelatable quantities in living systems.

**Fig. 2.**
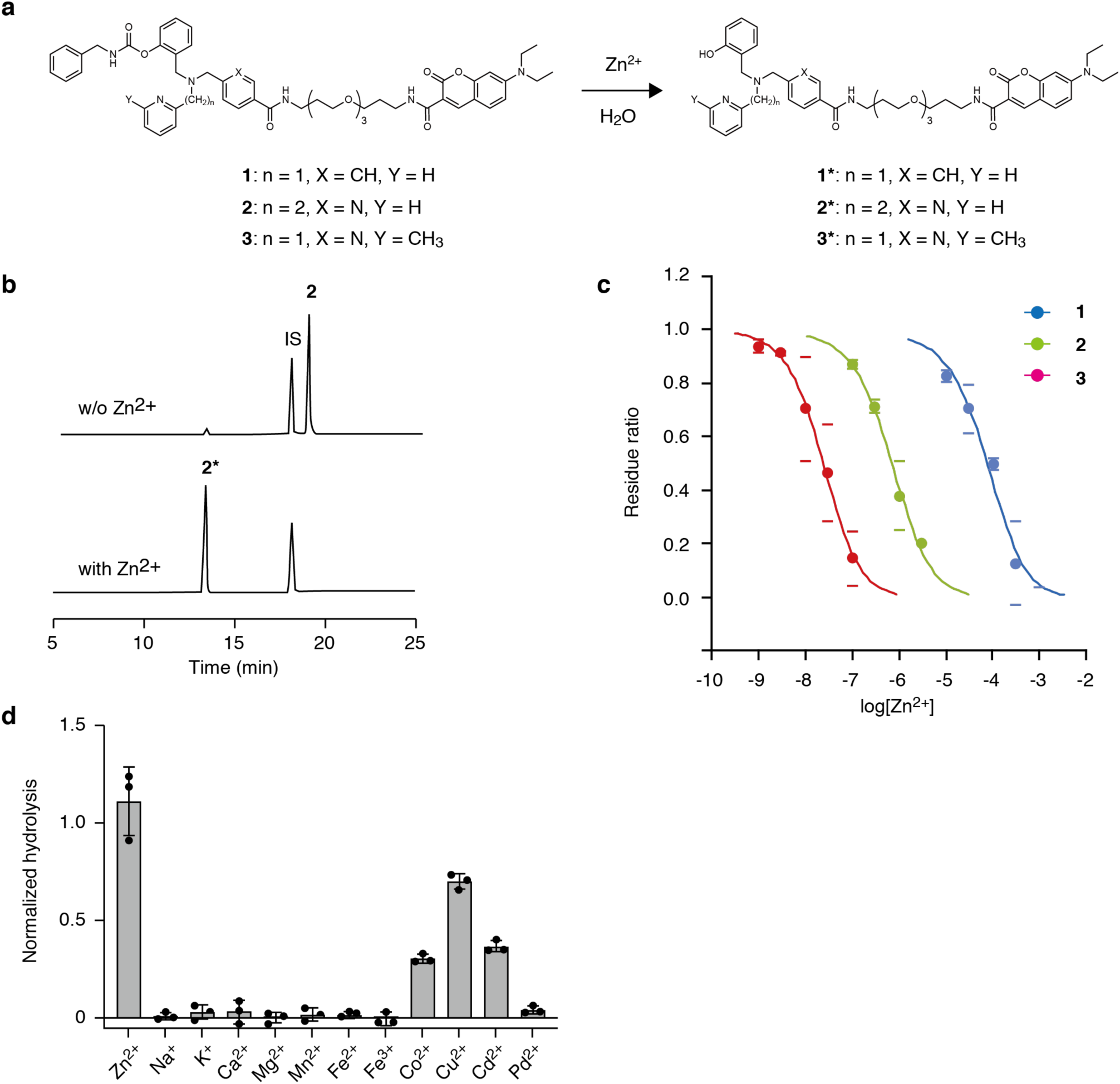
In vitro characterization of Zn^2+^-promoted hydrolysis of the carbamate linkage. **a**, Scheme of Zn^2+^-promoted hydrolysis of compounds **1**, **2** and **3**. **b**, HPLC chromatograms of **2** (1 µM) after incubation in 100 mM HEPES buffer (*I* = 0.1 (KOH + KCl), pH 7.4) at 37°C for 60 min in the presence and absence of Zn^2+^ (3 µM). IS, internal standard. **c**, Zn^2+^ concentration dependent hydrolysis of **1**, **2** and **3**. 1 µM of **1**, **2**, or **3** was incubated with varied concentrations of ZnSO_4_ at 37 °C for 60 min. Residue ratio = **1**/(**1** + **1\***), **2**/(**2** + **2\***) or **3**/(**3** + **3\***). Data are presented as mean values ± s.d. (n = 3). **d**, Metal ion selectivity of **2**. 5 mM of Na^+^, K^+^, Ca^2+^, Mg^2+^ or 1 µM of other metal ions were incubated with 1 µM of **2** in the buffer at 37 °C for 60 min. Normalized hydrolysis = **2\***/IS. Data are presented as mean values ± s.d. (n = 3).

### Construction of neuroreceptor-based fluorescent Zn^2+^ sensors on cultured cells

Having a set of Zn^2+^-responsive modules in hand, we then constructed neuroreceptor-based fluorescent Zn^2+^ sensors in cultured cells, using the AMPAR as a model receptor. In living HEK293T cells transiently expressing GluA2 (a subunit of AMPAR), LDAI chemistry allowed us to label GluA2 selectively using **CAM2 (TCO, Ax555)** for 4 h, followed by IEDDA click chemistry, that is, the addition of MeTz-tethered **Zinc probes** (30 min incubation), to prepare the semisynthetic GluA2-based Zn^2+^ sensor. The cell lysates were then analyzed by SDS-PAGE in-gel fluorescence imaging (Fig. 3a). Ax647 fluorescence derived from **Zinc probes** showed a single band corresponding to GluA2 (Fig. 3b–d), indicating selective anchoring of **Zinc probes** to AMPARs. To confirm Zn^2+^-induced cleavage of the probes, the cells were treated with exogenous Zn^2+^ after tethering the **Zinc probes**. A Zn^2+^-concentration-dependent decrease in the fluorescence band intensity was observed, which gave EC_50_ values of 3.0 × 10^−5^ M, 3.5 ×10^−7^ M and 2.3 × 10^−8^ M for **Zin(1), Zin(2)** and **Zin(3)**, respectively, being almost consistent with results obtained in vitro (Fig. 3e). Live-cell CLSM imaging showed that both Ax555 (**CAM2**) and Ax647 (**Zinc probes**) fluorescence localized to the cell surface (Fig. 3f–h). Upon Zn^2+^ treatment, the Ax647 fluorescence dramatically decreased, while the Ax555 signal remained essentially unchanged. Notably, ratiometric imaging (Ax555/Ax647) enabled quantitative analysis of Zn^2+^-induced cleavage of **Zinc probes**, normalizing for GluA2 expression and CAM2 anchoring, which more accurately reflect changes in Zn^2+^ concentrations. The GABA_A_R-based Zn^2+^ sensor was also constructed in living HEK293T cells using a similar protocol with **Fmz (TCO, Ax555)** as an anchoring reagent and **Zinc probes**, and we confirmed similar Zn^2+^-responsive ratiometric changes (Ax555/Ax647) in the peripheral region of HEK293T cells. (Supplementary Fig. 3). These results indicate that the receptor-based sensors using **Zinc probes** with varying affinities enable the detection of Zn^2+^ by CLSM imaging (Fig. 3i).

**Fig. 3.**
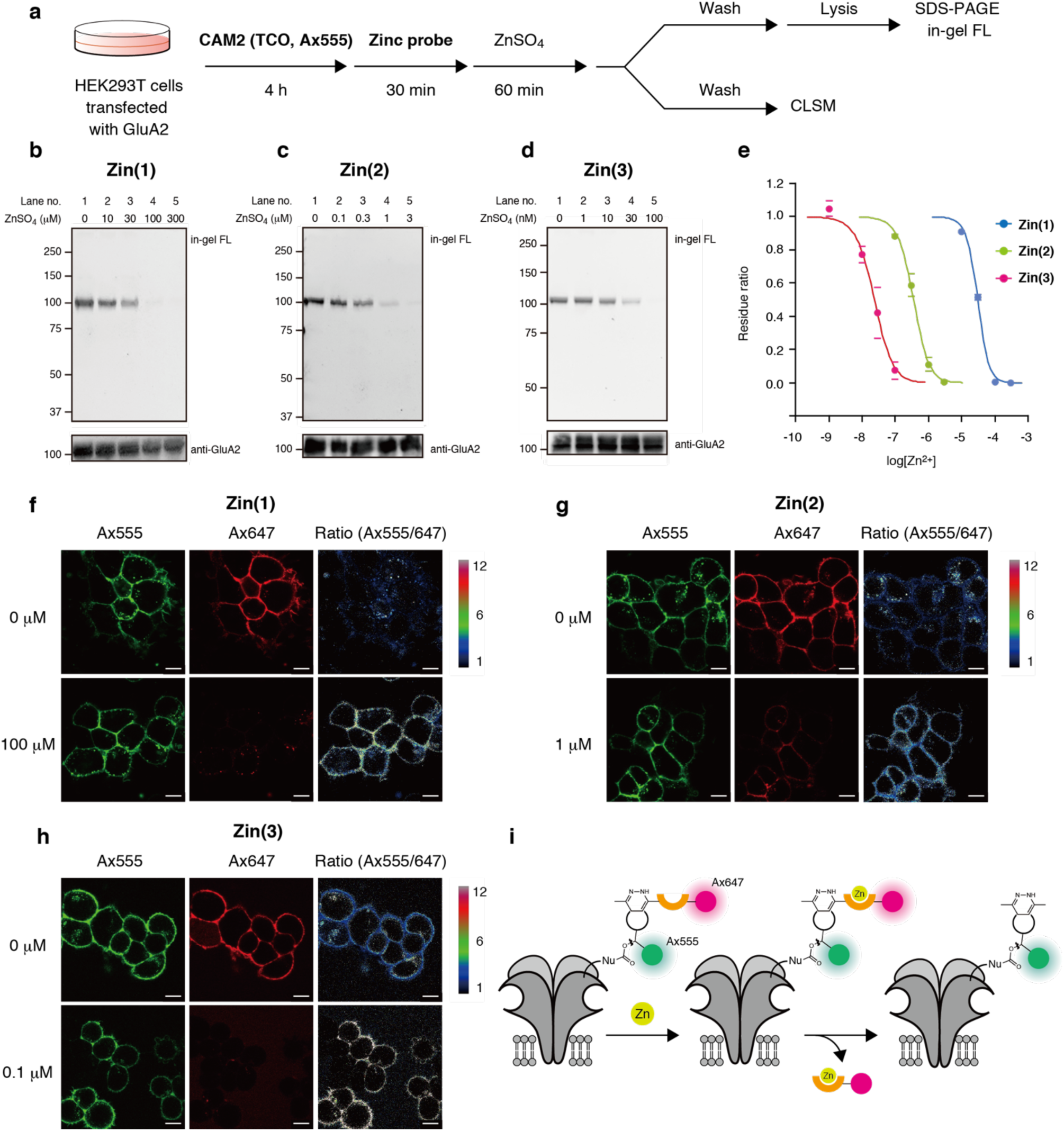
Construction of neuroreceptor-based fluorescent Zn^2+^ sensors on cultured cells. **a**, Experimental workflow in Fig. 3. HEK293T cells transiently expressing GluA2 were sequentially incubated with **CAM2 (TCO, Ax555)** (1 µM) for 4 h, followed by incubation with **Zin(1)**, **Zin(2)** or **Zin(3)** (1 µM) for 30 min. Subsequently, ZnSO_4_ was added into the medium at the indicated concentrations, and the cells were incubated for another 60 min. **b**–**d**, SDS-PAGE in-gel fluorescence and western blot analysis of lysates of HEK293T cells expressing AMPARs after labeling with **Zn(1)**, **Zin(2)** and **Zn(3)**, and their response to exogenously added Zn^2+^. **e**, Plots of concentration dependent cleavage of **Zn(1)**, **Zin(2)** and **Zn(3)** with Zn^2+^ in Fig. 3b–d. Data are presented as mean values ± s.d. (n = 3). **f–h**, CLSM imaging of HEK293T cells expressing AMPARs after labeling with **Zin(1)**, **Zin(2)** and **Zin(3)**. Cells were incubated for 60 min in the presence and absence of Zn^2+^ before CLSM imaging. Scale bar = 10 µm. **i**, Schematic illustration of neuroreceptor-based Zn^2+^ sensors.

### Imaging-based analyses of activity-dependent Zn^2+^ release at specific synapses in acute brain slices

To evaluate synaptic Zn^2+^ concentrations in the central nervous systems (CNS) using our neuroreceptor-based semisynthetic Zn^2+^ sensors, we applied them to acute hippocampal slices, as an ex vivo model of the CNS. Specifically, the **Zinc probes** were tethered to AMPARs to target excitatory glutamatergic synapses, where Zn^2+^ is accumulated in presynaptic vesicles together with glutamate and released to the synaptic cleft in an activity-dependent manner. In these experiments, we anchored TCO/Ax555 to AMPAR in the living brain using **CAM2 (TCO, Ax555)**, followed by **Zinc probe** tethering in the acutely prepared slice samples. In detail, **CAM2 (TCO, Ax555)** was directly injected into the lateral ventricle (LV) of a living mouse brain under anesthesia, followed by incubation for 24 h to label AMPARs selectively in vivo. Next, acute hippocampal slices were prepared from the excised brain and incubated for 15 min in artificial cerebrospinal fluid (ACSF) containing **Zinc probes** (Fig. 4a). Western blot analysis of the tissue homogenates using an anti-Ax647 antibody revealed a prominent band corresponding to GluA2, confirming the tethering of **Zinc probes** to AMPARs (Fig. 4b). CLSM imaging of the fixed slices showed strong colocalization between the Ax647 (**Zinc probes**) and the Ax555 (AMPAR) signals (Fig. 4c). Notably, densely distributed, bright puncta approximately 1 µm or less in diameter, were clearly visualized by high-resolution imaging of the hilus area in the dentate gyrus (DG), with a clear colocalization of the Ax647 and Ax555 signals (Fig. 4f). These results reveal the successful construction of fluorescent Zn^2+^ sensors at the AMPAR-locating glutamatergic synapses through the 2-step modification (brain/slice) protocol consisting of LDAI and click chemistry.

**Fig. 4.**
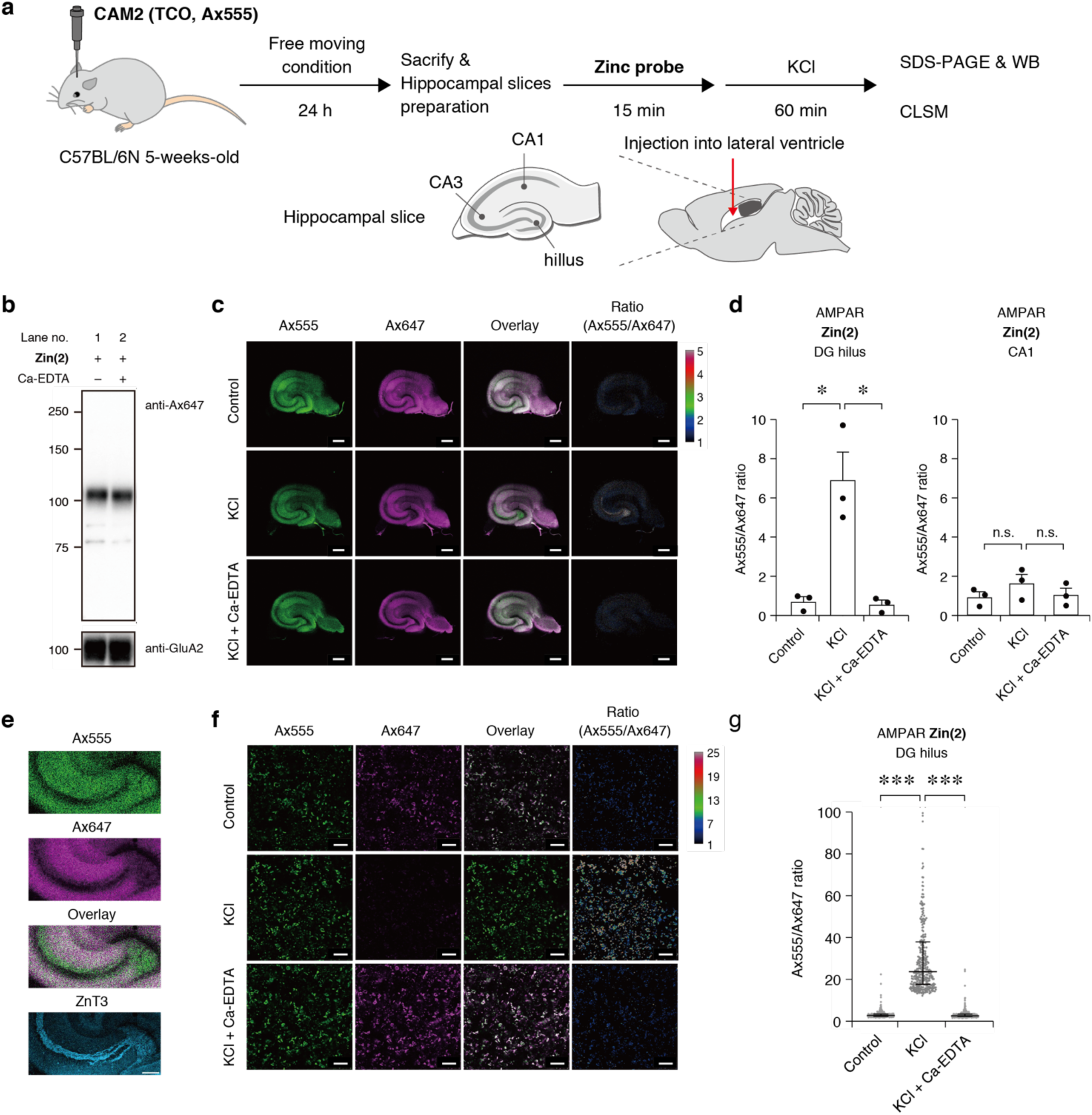
Recording of activity-dependent Zn^2+^ release at glutamatergic synapses in acute brain slices. **a**, Experimental workflow in Fig. 4. **CAM2 (TCO, Ax555)** (100 µM, 4.5 µL) was directly injected into the lateral ventricle (LV) of a living mouse brain, and 24 h after injection, the hippocampus was isolated. Subsequently, acute hippocampal slices (300 µm in thickness) were prepared and incubated with **Zin(2)** (1 µM) for 15 min. To induce the Zn^2+^ release, the slices were treated with 50 mM KCl for 60 min. **b**, Western blot analysis of labeling of **Zin(2)** to AMPARs. After incubation with **Zin(2)**, the slices were washed and further incubated in ACSF for 60 min. In lane 2, 10 mM Ca-EDTA was co-incubated with **Zin(2)**. **c**, CLSM imaging of the entire hippocampal slices labeled with **Zin(2)**. The slices were fixed with 4% PFA/PBS(–) containing 1 mM EDTA. Scale bar = 500 µm. **d**, The Ax555/Ax647 ratio values in the hilus of dentate gyrus (DG) and CA1 areas in Fig. 4c. Data represents mean values ± s.e.m. (n = 3). Student’s unpaired t-test with a two tailed distribution. *: *p* < 0.05. n.s.: not significant (*p* > 0.05). **e**, Enlarged CLSM images of hippocampal slices and colocalization analysis with anti-ZnT3 (rabbit, Almone Labs AZT-013, ×1000). Imaging was performed using a 5× objective (NA = 0.25). Scale bar = 200 µm. **f**, High-resolution confocal images of hilus of DG. Scale bar = 5 µm. **g**, Quantification of the Ax555/Ax647 ratios of puncta in Fig. 4f. The data represents mean values ± s.e.m. (n > 300). Student’s unpaired t-test with a two tailed distribution. ***: *p* < 0.001.

We next tested whether the Zn^2+^ sensors enabled visualization of synaptically released Zn^2+^ in response to KCl-evoked presynaptic depolarization. Slices containing the AMPAR-based Zn^2+^ sensors were incubated for 60 min in ACSF containing 50 mM KCl, followed by tissue fixation with paraformaldehyde (PFA). CLSM imaging of the entire hippocampal slice revealed a marked decrease in the Ax647 signal of **Zin(2)** in the mossy fiber regions, including the CA3 stratum lucidum and hilus of the DG. By contrast, Ax647 fluorescence in other hippocampal regions remained largely unchanged (Fig. 4d). The composite (Ax555 + Ax647) and ratiometric (Ax555/Ax647) images highlighted the selective hydrolysis of **Zin(2)** only in the mossy fiber region where the Zn^2+^ transporter (ZnT3) is highly expressed (Fig. 4e and Supplementary Fig. 4) and the labile Zn^2+^ may be condensed. Such changes in Ax647 of **Zin(2)** were not observed after co-incubation of KCl and Ca-EDTA, confirming that the KCl-induced and mossy fiber-specific fluorescence changes can be ascribed to the extracellular Zn^2+^ release. Furthermore, high-resolution imaging enabled us to observe Zn^2+^ release at the synaptic level. The punctate Ax647 signal from **Zin(2)** in the hilus of DG was significantly reduced by KCl-induced depolarization, resulting in a marked increase in the Ax555/Ax647 ratio (Fig. 4f,g). It is noteworthy that the size, morphology and intensity of the synaptic puncta visualized with Ax555 (AMPAR) remained unchanged, compared with those under the untreated conditions, implying the synaptic structures were preserved under the KCl stimulation.

Unlike excitatory glutamatergic synapses, inhibitory GABAergic synapses lack both ZnT3 and vesicular Zn^2+^ in their presynaptic terminals. To evaluate the local Zn^2+^ concentration at GABAergic synapses, we constructed the Zn^2+^ sensors on the GABA_A_R scaffold with a two-step (brain/slice) protocol as used for the AMPAR (Supplementary Fig. 5a). Western blot confirmed the selective tethering of **Zin(2)** to GABA_A_R (Supplementary Fig. 5b). CLSM imaging showed highly colocalized signals of **Zin(2)** with GABA_A_R (Supplementary Fig. 5c). Interestingly, KCl stimulation did not cause noticeable changes in either the Ax647 signal of **Zin(2)** or the Ax555/Ax647 ratio throughout the hippocampus, including in the mossy fiber region (Supplementary Fig. 5d). This suggests that the released Zn^2+^ into the glutamatergic synaptic cleft is rapidly diluted and/or recovered in hippocampal slice samples, thereby the local Zn^2+^ concentration in the proximity of GABAergic synapses is not elevated. Collectively, these results demonstrate that our neuroreceptor-based Zn^2+^ sensors in brain slice experiments can detect synaptically released Zn^2+^ at two representative synapse types: excitatory and inhibitory.

### Recording and quantitative analysis of synaptic Zn^2+^ in the living brains

Finally, we attempted to visualize and quantitatively analyze the activity-dependent synaptic Zn^2+^ release in the living brain using our neuroreceptor-based fluorescent Zn^2+^ sensors. Although acute brain slices are among the closest experimental models of the CNS, it would be ideal to study Zn^2+^ dynamics in the context of the living brain. According to our recent report,^30^ chemical KI enabled us to tether **Zinc probes** to a target neuroreceptor in the living mouse brain. After anchoring AMPARs or GABA_A_Rs with TCO and Ax555 by LV injection of **CAM2 (TCO, Ax555)** or **Fmz (TCO, Ax555)** (100 µM, 4.2 µL), respectively, a **Zinc probe** was subsequently injected into the LV site of the mouse brain (100 µM, 4.2 µL) (Fig. 5a). The hippocampus was isolated 3 h after the injection of **Zinc probe** and homogenates were analyzed by SDS-PAGE and in-gel fluorescence imaging. In the SDS-PAGE analysis of a **CAM2 (TCO, Ax555)**-injected hippocampal homogenate, a single Ax647-fluorescence band of approximately 100 kDa was observed, corresponding to the endogenous AMPAR subunit, indicating the selective modification of endogenous AMPARs with the **Zinc probe** (Fig. 5b). In the CLSM images of the whole brain cryosections, Ax647-fluorescence signals were observed from the several brain areas, such as hippocampus and molecular layer of cerebellum, where AMPARs are abundantly expressed (Fig. 5c). A zoomed-in image shows the strong fluorescence is derived from many puncta (Fig. 5f). These punctate signals were observed between the signals of anti-Homer (postsynaptic marker) and anti-Bassoon (presynaptic marker) antibodies, confirming the selective tethering of the **Zinc probe** to synaptic clefts. Similarly, both SDS-PAGE analysis of hippocampal homogenates and CLSM imaging of cryosections obtained from **Fmz (TCO, Ax555)**-injected mouse brains revealed the selective modification of GABA_A_Rs (Fig. 5d,e). Collectively, the results indicate that both AMPAR- and GABA_A_R-based Zn^2+^-sensors were successfully constructed in the living mouse brains through this 2-step (brain/brain) protocol. To confirm the stability of **Zinc probes**, we varied the incubation time from 3 to 6 or 9h after the **Zin(3)** injection, after which no further substantial changes in the Ax555/Ax647 ratio were observed in the DG hilus area of hippocampus (Supplementary Fig. 6). Given the EC_50_ value of **Zin(3)**, we conclude that the basal (extracellular) Zn^2+^ concentration at the glutamatergic synaptic clefts is below 20 nM.

**Fig. 5.**
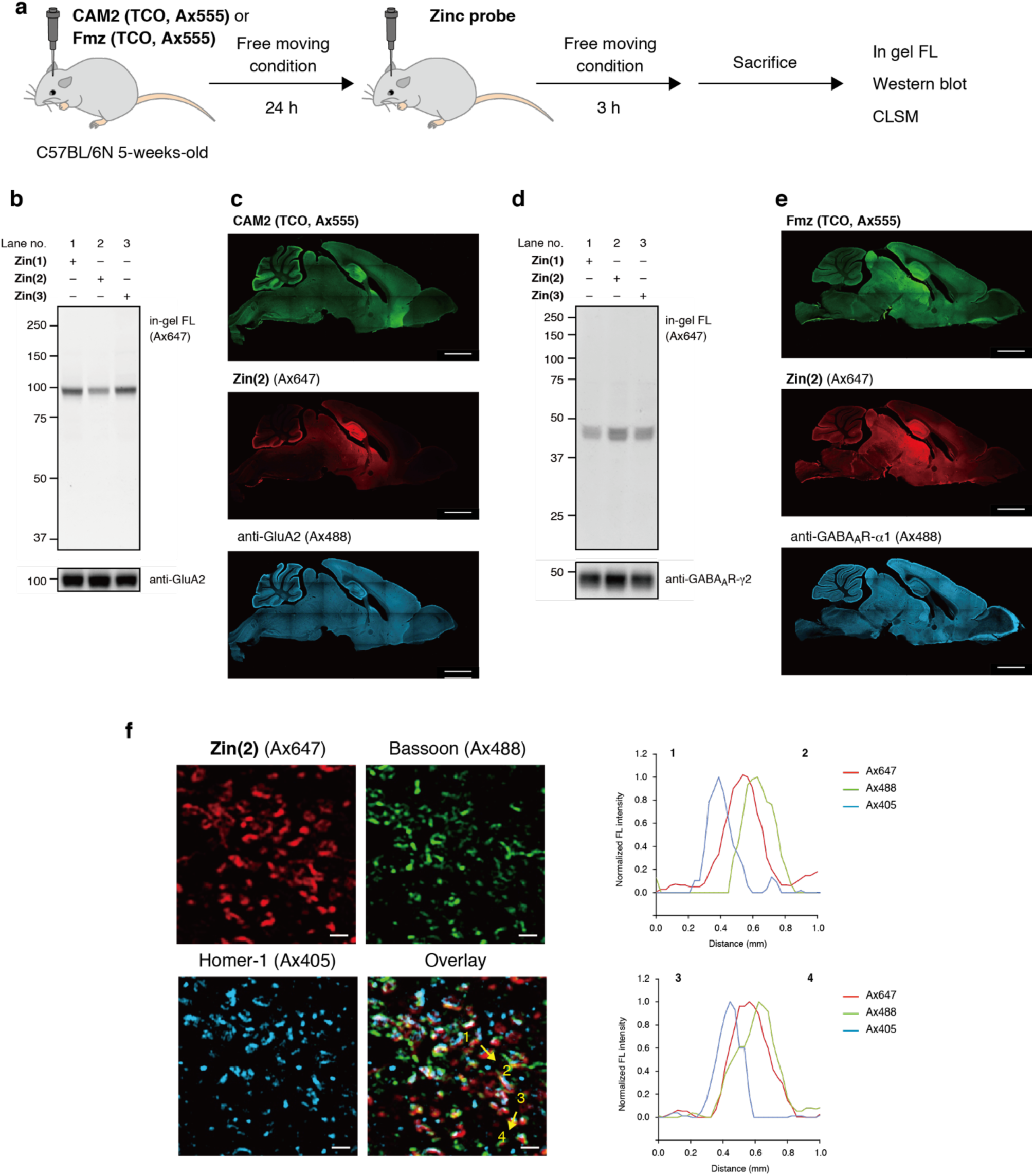
Construction of neuroreceptor-based Zn^2+^ sensors in a living mouse brain. **a**, Experimental workflow in Fig. 5. **CAM2 (TCO, Ax555)** (100 µM, 4.2 µL) or **Fmz (TCO, Ax555)** (100 µM, 4.2 µL) was directly injected into the lateral ventricle (LV) of a living mouse brain, and 24 h after injection, **Zin(1)**, **Zin(2)** or **Zin(3)** (100 µM, 4.2 µL) was injected into the LV. For SDS-PAGE and western blot analysis, 3 h after the **Zin probe** injection, hippocampus was isolated and a homogenate was prepared. For CLSM imaging, the mice were perfused transcardially with 4% PFA/PBS(–) containing 1 mM EDTA and whole brain cryosections were prepared by cryostat. CLSM imaging was conducted using a 5× (NA, 0.25) or 63× (NA, 1.4) objective. **b**, SDS-PAGE in-gel fluorescence and western blot analysis of **Zin(1)**, **Zin(2)** and **Zn(3)** labeling to AMPARs in a living mouse brain. **c**, CLSM imaging of the sagittal cryosection of a mouse brain labeled with **CAM2 (TCO, Ax555)** and **Zin(2)**. The cryosection was immunostained with anti-GluA2 antibody (rabbit, abcam ab206293, ×300). Scale bar, 2000 µm. **d**, SDS-PAGE in-gel fluorescence and western blot analysis of **Zin(1)**, **Zin(2)** and **Zn(3)** labeling to GABA_A_Rs. **e**, CLSM imaging of the sagittal cryosection of a mouse brain labeled with **Fmz (TCO, Ax555)** and **Zin(2)**. The cryosection was immunostained with anti-GABA_A_R-α1 antibody (mouse, Millipore 06-868, ×300). Scale bar = 2000 µm. **f**, High-resolution CLSM images of DG hilus area labeled with **CAM2 (TCO, Ax555)** and **Zin(2**). The mouse brain cryosection was immunostained with pre- and post-synapse markers (anti-Bassoon (rabbit abcam ab82958, ×1000) and anti-Hormer-1 (mouse, abcam ab184955, ×1000), respectively). 63× objective, Zeiss Airy scan mode. Fluorescence intensities of Alexa Fluor 405 (Ax405) (anti-Hormer-1), Alexa Fluor 488 (Ax488) (anti-Bassoon) and Ax647 (**Zin(2)**) were analyzed by line plots. Scale bar = 1 µm.

Using AMPAR-based fluorescent sensors constructed in the living brain, we sought to detect synaptic Zn^2^ release facilitated by kainic acid (KA)-induced seizures.^9^ Three hours after **Zinc probe** injection, the mice were injected intraperitoneally with KA or saline at a dose of 25 mg/kg body weight (Fig. 6a). We confirmed that the KA-treated mice exhibited clear epilepsy-like behavior within 1 h. Three hours after KA administration, the mice were perfused transcardially with 4% PFA/PBS(–) containing 1 mM EDTA under deep anesthesia with isoflurane. The CLSM imaging of cryosections of KA-treated mice showed a noticeable reduction in the Ax647 signals of **Zin(2)** probe (EC_50_ = 0.35 µM) in the DG hilus and CA3 areas of the hippocampus while retaining the Ax555 signals, resulting in an increase in the Ax555/Ax647 ratio (5.5-fold) compared with saline-injected controls (Fig. 6b,e). These findings suggest that KA-induced seizures increase Zn^2+^ release at glutamatergic synapses at the mossy fiber axon terminals. By contrast, in the case of **Zin(1)** (EC_50_ = 30 µM)-injected mice, a negligible difference was observed between KA- and saline-treated mice (Fig. 6c,e and Supplementary Fig. 7a). Based on the EC_50_ values of the two probes, we concluded that the local concentration of Zn^2+^ around synaptic AMPARs in DG hilus and CA3 areas is much higher than 0.35 µM, but does not reach 30 µM (Fig. 6i).

**Fig. 6.**
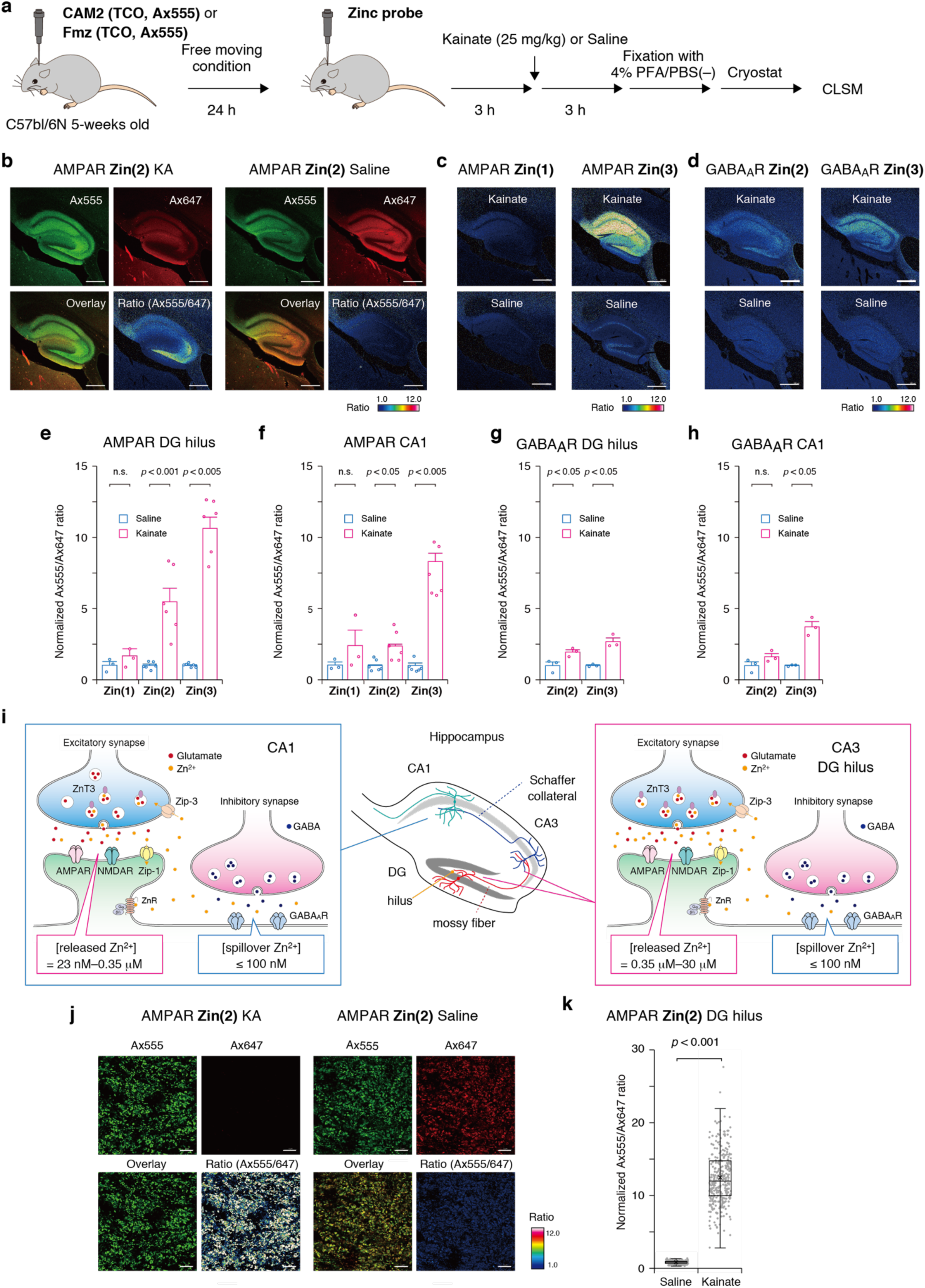
Recording of activity dependent synaptic Zn^2+^ release in living mouse brains. **a**, Experimental workflow in Fig. 6. AMPAR or GABA_A_R-based Zn^2+^ sensors were constructed as described in Fig. 5. Three hours after the injection of **Zin(1), Zin(2)** or **Zin(3)** into a lateral ventricle (LV) of a living mouse brain, the mice were injected intraperitoneally with KA (25 mg/kg mouse body weight) or saline. The mice were perfused transcardially with 4% PFA/PBS(–) containing 1 mM EDTA at 3 h after the KA or saline injection. CLSM imaging was conducted using a 5× (NA, 0.25) or 63× (NA, 1.4) objective. **b**, CLSM imaging of hippocampus area of mouse brain cryosections labeled with **CAM2 (TCO, Ax555)** and **Zin(2)**. scale bar = 500 µm. **c**, CLSM ratio imaging of hippocampus area of mouse brain cryosections labeled with **CAM2 (TCO, Ax555)** followed by the injection of **Zin(1)** or **Zin(3)**. scale bar = 500 µm. **d**, CLSM ratio imaging of hippocampus region of mouse brain cryosections labeled with **Fmz (TCO, Ax555)** followed by the injection of **Zin(2)** or **Zin(3)**. scale bar = 500 µm. **e**, Normalized Ax555/Ax647 ratio values in the DG hilus area of mouse brain cryosections labeled with **CAM2 (TCO, Ax555)** followed by the injection of **Zin(1), Zin(2)** or **Zin(3)**. The mean Ax555/Ax647 ratio value of saline-injected mouse under each condition is defined as 1. Data represents mean values ± s.e.m. (biological replicates, n = 3 or 6). Student’s unpaired t-test with a two tailed distribution. **f**, Normalized Ax555/Ax647 ratio values in the CA1 area of mouse brain cryosections labeled with **CAM2 (TCO, Ax555)** followed by the injection of **Zin(1), Zin(2)** or **Zin(3)**. The mean Ax555/Ax647 ratio value of saline-injected mouse under each condition is defined as 1. Data represents mean values ± s.e.m. (biological replicates, n = 3 or 6). Student’s unpaired t-test with a two tailed distribution. **g**, Normalized Ax555/Ax647 ratio values in the DG hilus area of mouse brain cryosections labeled with **Fmz (TCO, Ax555)** followed by the injection of **Zin(2)** or **Zin(3)**. The mean Ax555/Ax647 ratio value of saline-injected mice under each condition is defined as 1. Data represents mean values ± s.e.m. (biological replicates, n = 3). Student’s unpaired t-test with a two tailed distribution. **h**, Normalized Ax555/Ax647 ratio values in the CA1 region of mouse brain cryosections labeled with **Fmz (TCO, Ax555)** followed by the injection of **Zin(2)** or **Zin(3)**. The mean Ax555/Ax647 ratio value of saline-injected mice under each condition is defined as 1. Data represents mean values ± s.e.m. (biological replicates, n = 3). Student’s unpaired t-test with a two tailed distribution. **i**, Estimated Zn^2+^ concentrations released into the glutamatergic and GABAergic synaptic clefts at DG hilus/CA3 and CA1 areas. **j**, High-resolution CLSM images of the hilus area of hippocampal DG labeled with **CAM2 (TCO, Ax555)** and **Zin(2)**. The mice were treated with KA or saline 3 h before the perfusion with cold 4% PFA/PBS(–) containing 1 mM EDTA. scale bar = 5 µm. **k**, Boxplot of the Ax555/Ax647 ratio values of the fluorescence bright spots in Fig. 6j. The number of ROIs was 300 for each of the images (n = 300). The horizonal line and × within each box indicate the median and mean, respectively. Boxes show interquartile range (IQR). Whiskers show 1.5 × IQR.

The axon terminals of Schaffer collaterals, which project from the hippocampal CA3 to CA1, are also known to contain Zn^2+^-positive synaptic vesicles.^38^ Notably, a KA-elicited increase in the Ax555/Ax647 ratio in the CA1 of **Zin(2)**-injected mice (2.4-fold) was smaller than the changes observed in the DG hilus and CA3, and a larger change in the ratio (8.3-fold) was observed in the CA1 when **Zin(3)** probe was used (Fig. 6c,f and Supplementary Fig. 7b). These results suggest that Zn^2+^ release in CA1 is substantially lower than in the DG hilus and CA3, and the concentration in the synaptic cleft is estimated to be much higher than 23 nM and approximately 0.35 µM based on the EC_50_ values of the probes (Fig. 6i).

KA-induced Zn^2+^ dynamics measured with GABA_A_R-based fluorescent sensors in the hippocampal DG hilus and CA1 showed smaller Ax555/Ax647 changes for **Zin(2)** and **Zin(3)** probes than for the corresponding AMPAR-based sensors (Fig. 6d,g,h and Supplementary Fig. 8). These findings suggest that KA-induced seizures evoke a Zn^2+^ spillover from excitatory to GABAergic synapses, where the Zn^2+^ concentrations near GABA_A_Rs are lower than those near AMPARs. From the EC_50_ values of the **Zin(2)** and **Zin(3)** probes (0.35 µM and 23 nM, respectively), the Zn^2+^concentrations spilled over into the inhibitory synapses were estimated to be ≤100 nM.

High-magnification CLSM images resolved Zn^2+^ release events at synaptic spatial resolution. In the hilus of the hippocampal DG of mice injected with the **Zin(2)** probe, fluorescence signals from Ax555 and Ax647 appeared as bright puncta of ≤1 µm in diameter that were clearly colocalized (Fig. 6j). KA-treatment reduced Ax647-fluorescent puncta, whereas Ax555 signals were retained, increasing the Ax555/Ax647 ratio in many synapses (Fig. 6k). Similar changes occurred with **Zin(3)**, whereas **Zin(1)** produced no detectable change in the Ax555/Ax647 ratio, even at synaptic-level spatial resolution (Supplementary Fig. 9). These CLSM data demonstrate that our neuroreceptor-based sensors can report Zn^2+^ dynamics near AMPARs in the excitatory hippocampal synapses in the living mouse brain with synaptic spatial resolution.

## Discussion

Chemical and genetically encoded fluorescent Zn^2+^ sensors have advanced our understanding of labile Zn^2+^in the CNS. However, these methods have still suffered from limited spatial resolution and insufficient quantitative capability, leaving the dynamics and functional roles of labile Zn^2+^ at the synaptic level unresolved. For example, reported concentrations of Zn^2+^ released into the synaptic cleft span a wide range (1–300 µM), and the true values have remained elusive.^39,40^ Although Zn^2+^ spillover into inhibitory synapses has been proposed, quantitative analyses appear lacking. Our chemical KI strategy enabled the conversion of endogenous AMPAR and GABA_A_R receptors into Zn^2+^ sensors in the living mouse brain, allowing us to record Zn^2+^ dynamics in both excitatory and inhibitory synaptic clefts. The bond-cleavage-based recording preserved the history of Zn^2+^ release, enabling in-depth, imaging-based analysis in fixed mouse brain cryosections. Using this approach, we quantified Zn^2+^ release at single-synapse resolution and quantitatively determined the concentrations of Zn^2+^ released into the glutamatergic synaptic clefts in several hippocampal regions. KA-induced seizure increased synaptic Zn^2+^ concentrations by more than an order of magnitude, from less than 20 nM (basal state) to higher than 0.35 µM (but not approaching 300 µM). By contrast, Zn^2+^ concentrations near GABA_A_Rs remained ≤100 nM, substantially lower than those near AMPARs, suggesting efficient labile Zn^2+^ clearance systems around excitatory synapses in the hippocampus. Moreover, KA-induced synaptic Zn^2+^ concentration elevations were markedly greater in the DG hilus/CA3 than in CA1. This regional difference is consistent with previous Timm staining analysis^8^ as well as with reports that nearly all glutamatergic vesicles in mossy fiber presynaptic terminals contain Zn^2+^, whereas only approximately 50% vesicles in Schaffer collateral terminals do so.^38^

Zn^2+^ co-released with glutamate may interact with multiple neuroreceptors, including NMDARs, AMPARs and GABA_A_Rs, and modulate their function. Previous reports using chelators for masking the mobile Zn^2+^ in the acute slices^5,6^ or in vivo^41^ have shown that Zn^2+^ act as a functional modulator of these neuroreceptors, most often producing inhibitory effects, although potentiation of AMPARs has also been reported. However, the synaptic Zn^2+^ concentration has been inferred only indirectly or quantified poorly. For NMDARs, which are tetramers composed of two GluN1 and two GluN2 subunits, measurements in vitro show that GluN2A-containing NMDARs have nanomolar affinity for Zn^2+^, whereas GluN1/GluN2B heteromers bind Zn^2+^ only in the tens-of-micromolar range.^42^ Given the KA-induced Zn^2+^ concentrations we determined in excitatory synaptic clefts, the elevated concentrations are sufficiently high enough to inhibit the GluN2A-containing NMDARs, but may remain below the threshold need to modulate GluN1/GluN2B heteromers.^43^ Reports on AMPARs regulation by Zn^2+^ are mixed, suggesting synapse-dependent or Zn^2+^ concentration-dependent modes of action.^6,41^ Yet the reported IC_50_ values for Zn^2+^ inhibition of AMPARs expressed in HEK293T cells (460–1750 µM) are substantially higher than the synaptic Zn^2+^ concentration we measured (<30 µM in all cases), indicating that direct Zn^2+^–AMPARs interactions are unlikely to contribute substantially under our experimental conditions.^44^ Zn²⁺ released from ZnT3-positive glutamatergic synapses can spill over to neighboring GABAergic synapses and influence long-term potentiation (LTP).^45^ GABA_A_Rs comprise complex pentameric assemblies with diverse combinations of subunits, and receptors containing the synapse-selective γ-subunit have IC_50_ values for Zn^2+^ inhibition of approximately 300 µM.^46^ Because our anchoring reagent selectively labels γ-subunit containing GABA_A_Rs,^29^ and because the Zn^2+^ concentrations we measured near these receptors after spillover are ≤100 nM, inhibition of synaptic GABA_A_Rs by Zn^2+^ is unlikely, even in the KA-triggered seizures. By contrast, GABA_A_Rs consist of α- and β-subunits have much lower IC_50_ values for Zn^2+^ (IC_50_ = 88 nM) and these are extrasynaptic pentamers. Therefore, Zn^2+^ leaked from excitatory synapses may preferentially inhibit the extrasynaptic GABA_A_Rs rather than γ-subunit containing synaptic receptors.^7,46^

Because our chemical KI can apply to the whole brain in living mice, application of our neuroreceptor-based Zn^2+^-sensors could be extended to target additional brain regions, such as the auditory cortex, amygdala, dorsal cochlear nucleus and olfactory bulb, to enable more quantitative and comprehensive analysis of Zn^2+^ functions in diverse neural circuits. Furthermore, the present chemical design may be adaptable to other biologically relevant metal ions whose roles in brain function remain to be elucidated.

## Methods

### Instruments for biochemical and biological experiments

SDS-polyacrylamide gel electrophoresis (SDS-PAGE) and western blotting were carried out using a Bio-Rad Mini-Protean III electrophoresis apparatus. UV-Vis spectra were measured using Shimazu UV-2600 UV-Vis spectrophotometer. Chemical luminescence signals generated with ECL Prime (Cytiva, Amersham) and fluorescence gel images were acquired with a FUSION-FX7 imaging system (Vilber-Lourmat). Cell and tissue imaging was performed with a confocal laser scanning microscope (CLSM) (Carl Zeiss LSM800) equipped with a 5× objective lens (numerical aperture (NA) = 0.25, dry objective), 63× objective lens (NA = 1.40, oil objective), and GaAsP detector. The excitation laser was derived from a 488 nm, 561 nm, 640 nm diode laser and was set to an appropriate wavelength depending on the dye. Airyscan mode (Carl Zeiss) was used in synapse-resolution imaging.

### HPLC analysis of Zn^2+^-promoted hydrolysis of the carbamate linkage in vitro

1 µM of **1**, **2** or **3** was incubated with 1 mM of EDTA or ZnSO_4_ at the indicated concentration in 100 mM HEPES buffer (pH 7.4, ionic strength, *I* = 0.1 (KOH/KCl)) at 37 °C for the indicated time. For **3**, free Zn^2+^ concentration was controlled by using 0–9.75 mM ZnSO_4_/10 mM nitrilotriacetic acid (NTA).^16^ 1 mM of EDTA was added to halt the Zn^2+^-promoted hydrolysis reaction. 7-(Diethylamino)coumarin-3-carboxylic acid (Tokyo Chemical Industry (TCI)) (2 µM) was added as an internal standard (IS). The reaction mixture was subjected to RP-HPLC analyses (column: YMC-Triart C18, TA12S05-2546WT, 250 × 4.6 mm; mobile phase: CH_3_CN (0.1% TFA)/H_2_O (0.1% TFA) = 30/70 to 70/30 (linear gradient over 30 min); flow rate: 1.0 mL/min; UV detection: 420 nm). The hydrolysis of **1**, **2** or **3** was quantified by residue ratio (**1**/(**1** + **1**\*), **2**/(**2** + **2\***) or **3**/(**3** + **3\***)). To evaluate the metal ion selectivity of **2**, 5 mM of NaCl, KCl, CaCl_2_, MgCl_2_ or 1 µM of ZnSO_4_, MnSO_4_, FeCl_2_·4H_2_O, FeCl_3_·6H_2_O, CoSO_4_·7H_2_O, CuSO_4_·5H_2_O, CdSO_4_·H_2_O, Pb(NO_3_)_2_ were incubated with 1 µM of **2** at 37 °C for 60 min. The hydrolysis of **2** was normalized relative to IS. Normalized hydrolysis = **2\***/IS. The EC_50_ values were determined by fitting the plots of residual ratio with a theoretical logistic equation.

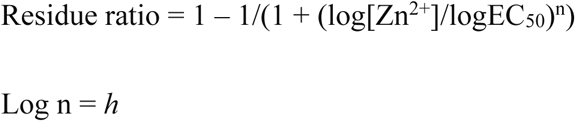

where *h* is a Hill’s coefficient.

### Cell culture

HEK293T cells (ATCC) were cultured in Dulbecco’s Modified Eagle Medium (DMEM)-GlutaMAX (Gibco) supplemented with 10% dialyzed fetal bovine serum (FBS) and 1% Antibiotic-Antimycotic (Anti-Anti) (Fujifilm) and incubated in a 5% CO_2_ humidified chamber at 37°C. Transient expression of GluA2^flip^(Q)^47^ or GABA_A_R-α1β3γ2^48^ was carried out with Lipofectamine 2000 (Invitrogen) according to the manufacturer’s instructions.

### Construction of receptor-based fluorescent Zn^2+^ sensors on HEK293T cells

For the chemical labeling of AMPAR or GABA_A_R with a **LDAI anchoring reagent**, HEK293T cells transfected with GluA2 or GABA_A_R-α1β3γ2 were incubated with 1 µM of **CAM2 (TCO, Ax555)** or **Fmz (TCO, Ax555)** in serum-free DMEM-GlutaMAX containing HEPES (Gibco) at 17 °C for 4 h. For the labeling with **Zinc probes**, the **anchoring reagent** solution was removed, and the cells were washed with HBS (20 mM HEPES, 107 mM NaCl, 6 mM KCl, 11.5 mM glucose, 2 mM CaCl_2_, 1.2 mM MgSO_4_, pH 7.4) (×1). The cells were treated with 1 µM **Zinc probes** in HBS at room temperature for 30 min. To quench the IEDDA reaction, 20 µM TCO-OH in HBS was then added and incubated for 5 min at room temperature. For **Zinc 3**, the second step labeling was conducted in the presence of 10 mM EDTA. To evaluate the Zn^2+^ response of the constructed sensors, the cells were treated with ZnSO_4_ at the indicated concentrations in HBS and incubated at room temperature for 1 h. For **Zinc 3**, 0–9.75 mM ZnSO_4_/10 mM NTA in 100 mM HEPES (pH 7.4, ionic strength, *I* = 0.1 (KOH/KCl)) were used to control free Zn^2+^ concentrations.

### SDS-PAGE and western blot analysis in HEK293T cells

The cells were washed three times with PBS(–) and lysed on ice with RIPA buffer (25 mM Tris–HCl, 150 mM NaCl, 1% Nonidet P-40, 0.1% sodium dodecyl sulfate, 0.25% sodium deoxycholate, pH 7.4) containing 1% protease inhibitor cocktail set III (Calbiochem, 39134) and 1 mM EDTA. The lysates were clarified by centrifugation at 13500 rpm for 10 min at 4 °C, and the protein concentrations were measured by BCA assay. After uniformizing the protein concentrations between samples, the lysates were then mixed with 5× Laemmli buffer (325 mM Tris-HCl, 15% SDS, 20% sucrose, and 0.2% bromophenol blue, pH 6.8) containing 250 mM of DTT. After shaking at room temperature for 1 h, the samples were applied to a 7.5% (for AMPARs) or 10% (for GABA_A_Rs) SDS-PAGE gel and imaged by in-gel fluorescence. For the western blot analysis, the proteins separated by SDS-PAGE were transferred onto Immun-Blot PVDF membranes (Bio-Rad). The membranes were blocked with 5% skim milk in Tris-buffered saline containing 0.05% Tween 20 (TBS-T) at room temperature for 1 h and incubated with primary antibodies at 4 °C overnight. After washing twice with TBS-T, the membranes were incubated with secondary antibodies at room temperature for 1 h. The following primary antibodies were used: rabbit anti-GluA2 (Abcam ab206293, ×2500); rabbit anti-GABA_A_R-γ2 (Synaptic Systems 224003, ×2000). The following secondary antibodies were used: goat anti-rabbit IgG-HRP conjugate (Cell signaling Technology 7074S, ×2000).

### CLSM imaging in HEK293T cells

HEK293T cells transfected with GluA2 or GABAAR-α1β3γ2 were seeded in 35 mm glass-bottom dishes (Matsunami D115304H) pre-coated with poly-L-lysine (Sigma-Aldrich). After constructing GluA2- or GABA_A_R-based fluorescent Zn^2+^ sensors, the cells were incubated with ZnSO_4_ as mentioned above. The cells were washed with HBS twice and subjected to CLSM imaging.

### Animal experiments

C57BL/6N mice (male, 5 weeks old, body weight of 18–23 g) were purchased from Japan SLC, Inc (Shizuoka, Japan). The animals were housed in a controlled environment (23 °C, 40–60% humidity, 12 h light–dark cycle) and had free access to food and water, according to the regulations of the Guidance for Proper Conduct of Animal Experiments by the Ministry of Education, Culture, Sports, Science and Technology of Japan. All experimental procedures were performed in accordance with the National Institute of Health Guide for the Care and Use of Laboratory Animals and were approved by the Institutional Animal Use Committees of Kyoto University.

### Construction of AMPAR-anchored fluorescent Zn^2+^ sensors in acute hippocampal slices

Under the deep anesthesia, 4.5 µL of **CAM2 (TCO, Ax555)** or **Fmz (TCO, Ax555)** (100 µM) in phosphate-buffered saline (pH 7.4) was directly injected into the right lateral ventricle (LV) (anteroposterior (AP) of -1.3 mm from bregma, mediolateral (ML) of +2 mm, depth of 2 mm) using a microinjector (Nanoliter 2010, World Precision Instruments) (600 nL/min). At 24 h after the injection, the mouse was sacrificed under the deep anesthesia with isoflurane and acute hippocampal slices (300 µm thickness) were prepared in an ice cold cutting solution (120 mM choline chloride, 3 mM KCl, 8 mM MgCl_2_, 1.25 mM NaH_2_PO_4_, 28 mM NaHCO_3_, 22 mM glucose, 0.5 mM L(+)-ascorbic acid) by bubbling with 95% O_2_. After recovery at 34 °C under 95% O_2_ for 1 h, the slices were incubated with 1 µM of **Zinc probe** in ACSF solution (125 mM NaCl, 2.5 mM KCl, 2 mM CaCl_2_, 1 mM MgCl_2_, 26 mM NaHCO_3_, 1.25 mM NaH_2_PO4, 10 mM glucose) at room temperature under 95% O_2_ for 15 min. For the visualization of synaptically released Zn^2+^, the slices were then washed twice with ACSF solution and incubated with 50 mM KCl for 60 min to induce depolarization. As a control, 10 mM Ca-EDTA was included during the incubation with **Zinc probes** and KCl to chelate extracellular Zn^2+^.

### SDS-PAGE and western blot analysis in acute hippocampal slices

SDS-PAGE and western blot analysis were conducted similarly as described in “SDS-PAGE and western blot analysis in HEK293T cells”. The following primary antibodies were used: mouse anti-Alx647 (Immunology Consulting Laboratory (ICL) M647-65A-400, ×2500); rabbit anti-GluA2 (Abcam ab206293, ×5000); rabbit anti-GABA_A_R-γ2 (Synaptic Systems 224003, ×4000). The following secondary antibodies were used: goat anti-mouse IgG-HRP conjugate (Cell Signaling Technology 7076S, ×2500) and goat anti-rabbit IgG-HRP conjugate (Cell Signaling Technology 7074S, ×2500).

### CLSM imaging in acute hippocampal slices

To prevent undesired metal-induced hydrolysis of **Zinc probes** during subsequent procedures, 1 mM EDTA was included in all solutions used for washing, fixation, permeabilization, and immunostaining. The slices were fixed with 4% paraformaldehyde (PFA) in PBS(–) at 4 °C overnight. For immunostaining, the slices were then permeabilized in 0.1% Triton X-100 in PBS(–) at room temperature for 15 min. After blocking in PBS(–) containing 2% bovine serum albumin (BSA), 2% normal goat serum (NGS) and 0.1% Triton X-100, at room temperature for 30 min, the slices were incubated with primary antibodies in PBS(–) containing 10% NGS and 0.1% Triton X-100 at 4 °C overnight. Subsequently, the slices were incubated with secondary antibodies in PBS(–) containing 10% NGS and 0.1% Triton X-100 at room temperature for 1 h. The following primary antibodies were used: rabbit anti-ZnT3 (Almone Labs AZT-013, ×1000). The following secondary antibodies were used: goat anti-rabbit IgG-Alexa488 conjugate (Invitrogen, A11008, ×1000).

### Construction of receptor-based fluorescent Zn^2+^ sensors in a living mouse brain

Under the deep anesthesia, 4.2 µL of **LDAI anchoring reagent** (100 µM) in PBS(–) (pH 7.4) was directly injected into the right LV (AP of -1.3 mm from bregma, ML of +2 mm, depth of 2 mm) using a microinjector (600 nl/min). At 24 h after the injection, **Zinc probe** (100 µM, 4.2 µL) was injected into the right LV. For western blot analysis, at 3 h after the injection, mice were sacrificed under the deep anesthesia with isoflurane and hippocampus was isolated. The isolated brain tissue was lysed in a RIPA buffer containing 1% protease inhibitor cocktail set III and 1 mM EDTA and gently mixed with a rotator at 4 °C for 1 h. The homogenates were centrifuged at 4 °C and 17700 × g for 10 min to remove the insoluble matter. SDS-PAGE and western blot analysis were conducted similarly as described in “SDS-PAGE and Western blot analysis in HEK293T cells”. The following primary antibodies were used: rabbit anti-GluA2 (Abcam ab206293, ×4000); rabbit anti-GABA_A_R-γ2 (Synaptic Systems 224003, ×4000). The following secondary antibody was used: goat anti-rabbit IgG-HRP conjugate (Cell Signaling Technology 7074S, ×4000). For CLSM imaging, at 3 h after the injection, mice were sacrificed under the deep anesthesia with isoflurane by transcardial perfusion with ice cold 4% PFA/PBS(–) containing 1 mM EDTA (pH 7.4). The isolated mouse brain samples were fixed overnight at 4 °C in 4% PFA/PBS(–) containing 1 mM EDTA. After washing with PBS(–) containing 1 mM EDTA (×3), the brain samples were immersed in 30% sucrose/PBS(–) containing 1 mM EDTA. Mouse brain cryosections were prepared using a cryostat (Leica, CM-1950). The following primary antibodies were used for immunostaining of mouse brain cryosections: rabbit anti-GluA2 (abcam ab206293, ×300), rabbit anti-GABA_A_R-α1 (Merck 06-868, ×300), mouse anti-Homer1 (abcam ab184955, ×1000) and rabbit anti-Bassoon (abcam ab82958, ×1000). The following secondary antibody was used: goat anti-rabbit IgG H&L (Alexa Fluor 488 conjugate) (Invitrogen, A32731, ×300) and goat anti-mouse IgG H&L (Alexa Fluor 405 conjugate) (Invitrogen, A31553, ×300).

### Synaptic zinc release in hippocampus of living mouse brain under kainite-induced seizures

Under the deep anesthesia, 4.2 µL of **LDAI anchoring reagent** (100 µM) in PBS(–) (pH 7.4) was directly injected into the right LV using a microinjector (600 nL/min). At 24 h after the injection, **Zinc probe** (100 µM, 4.2 µL) was injected into the LV. At 3 h after the injection, the mice were injected intraperitoneally with KA (25 mg/kg body weight) or saline. At 3 h after KA-administration, the mice were sacrificed under the deep anesthesia with isoflurane by transcardial perfusion with ice cold 4% PFA/PBS(–) containing 1 mM EDTA (pH 7.4). The isolated mouse brain samples were fixed overnight at 4 °C in 4% PFA/PBS(–) containing 1 mM EDTA (pH 7.4). After washing with PBS(–) containing 1 mM EDTA (×3), the brain samples were immersed in 30% sucrose/PBS(–) containing 1 mM EDTA. Brain slices were prepared using a cryostat (Leica, CM-1950).

### Data availability

Detailed synthetic procedures and compound characterization are available in the Supplementary Information.

## Acknowledgements

The authors thank T. Gonda, Y. Nabeta, K. Nishizawa and M. Ishikawa for technical support of biological experiments. The authors also thank Robin James Storer, PhD, from Edanz (https://jp.edanz.com/ac) for editing a draft of this manuscript. This work was supported by MEXT/JSPS KAKENHI Grant-in-Aid for Specially Promoted Research (grant number 23H05405 to I.H.), MEXT/JSPS KAKENHI Grant-in-Aid for Scientific Research (B) (grant number 24K01627 to H.N.), MEXT/JSPS KAKENHI Grant-in-Aid for Scientific Research (C) (grant number 23K04960 to S.S.).

## Author contributions

H.Z., H.N. and I.H. initiated and designed the project. H.Z., S.S., K. Nakamura, Q.C., K. Nakajima and H.N. performed synthesis, compounds characterization and chemical labeling experiments in HEK293T cells and brains. H.Z. S.S., H.N. and I.H. prepared the manuscript with contributions from the other authors.

