## Supplementary Information for "Quantitative analysis of synaptic zinc in the brain by covalent chemistry-based semisynthetic biosensors"

**Table of contents**

**Supplementary Figures** 3

**General materials and methods for organic synthesis** 12

**Synthesis of 1** 14

**Synthesis of 2** 17

**Synthesis of 3** 20

**Synthesis of Zin(1)** 23

**Synthesis of Zin(2)** 26

**Synthesis of Zin(3)** 29

**Synthesis of Fmz (TCO, Ax555)** 31

**Supplementary References** 33

**Supplementary Figures**

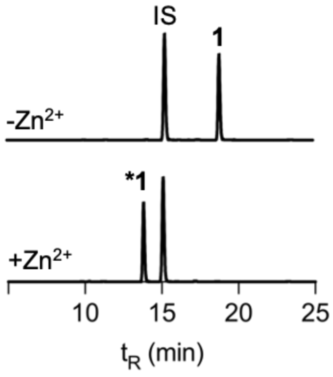


**Supplementary Fig. 1 | HPLC chromatograms of 1 in the absence and presence of $Zn^{2+}$.** IS: internal standard. 1 μM of **1** was incubated with 1 mM of EDTA (–$Zn^{2+}$) or 1 mM of $ZnSO_4$ (+$Zn^{2+}$) in 100 mM HEPES buffer (pH 7.4, ionic strength, $I$ = 0.1 (KOH/KCl)) at 37 ºC for 60 min.

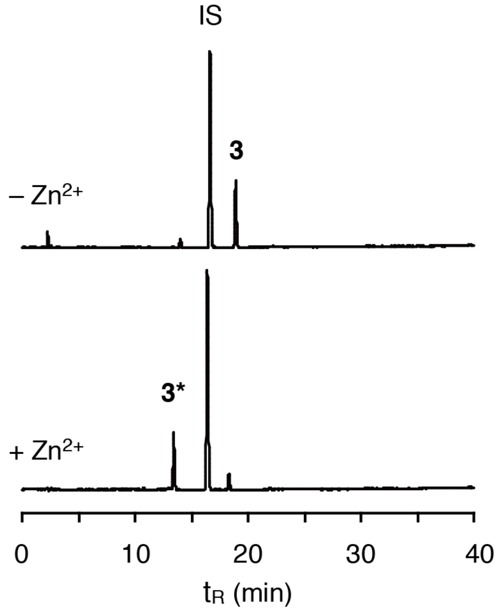


**Supplementary Fig. 2 | HPLC chromatograms of 3 in the absence and presence of $Zn^{2+}$.** IS: internal standard. 1 μM of **3** was incubated with 1 mM of EDTA (–$Zn^{2+}$) or 0.1 μM of $ZnSO_4$ (+$Zn^{2+}$) in 100 mM HEPES buffer (pH 7.4, ionic strength, $I$ = 0.1 (KOH/KCl)) at 37 ºC for 60 min.

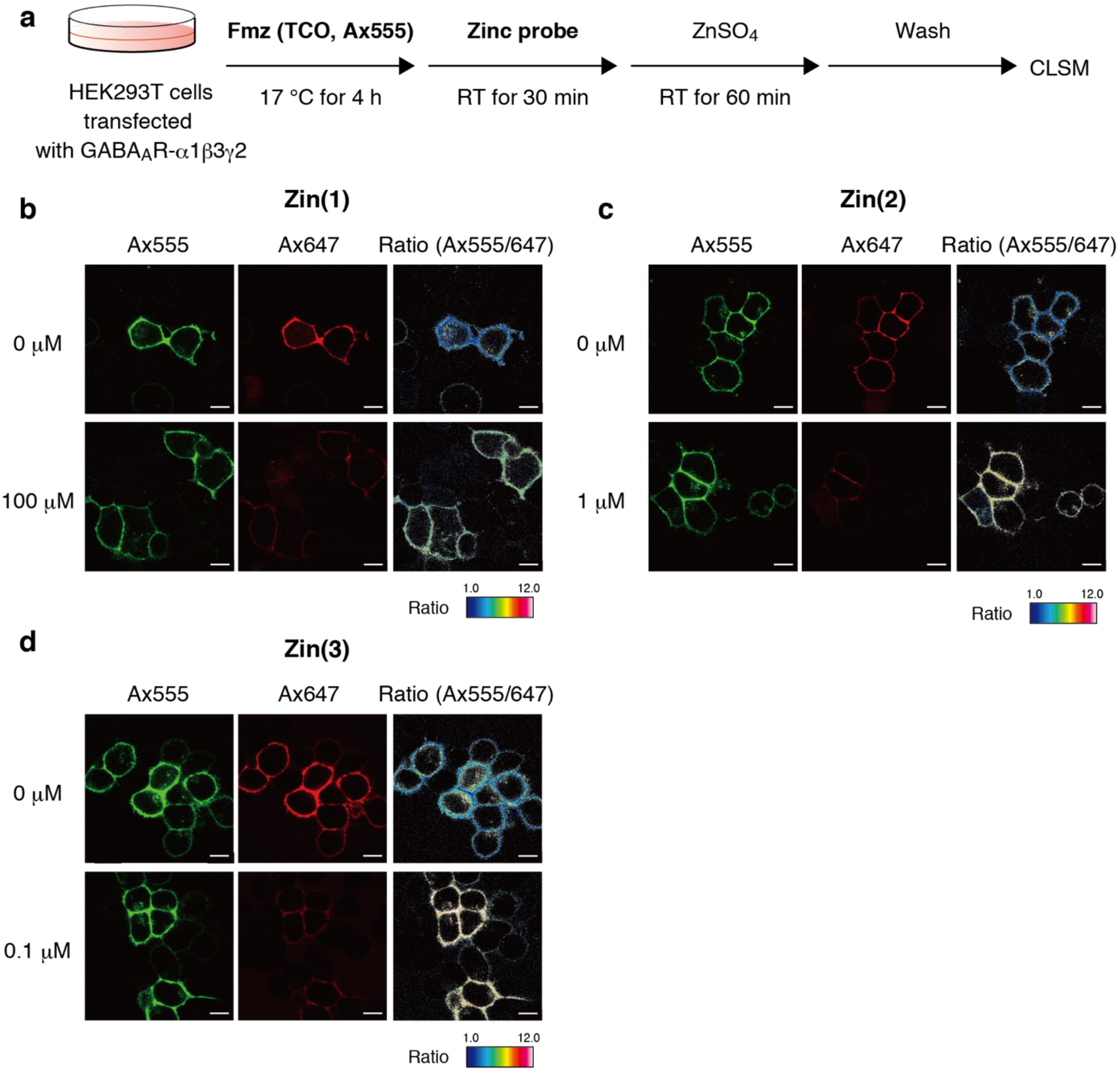


**Supplementary Fig. 3 | Construction of GABA$_A$R-based fluorescent Zn$^{2+}$ sensors on cultured HEK293T cells. a**, Experimental workflow in Supplementary Fig. 3. HEK293T cells transiently expressing GABA$_A$R-α1β3γ2 were sequentially incubated with **Fmz (TCO, Ax555)** (1 μM) at 17 °C for 4 h, followed by **Zn(1)**, **Zn(2)** or **Zin(3)** (1 μM) at room temperature for 30 min. Subsequently, $ZnSO_4$ was added at the indicated concentrations, and the cells were incubated for another 60 min at room temperature. **b–d**, CLSM images of HEK293T cells expressing GABA$_A$R-α1β3γ2 after labeling with **Zn(1)** (**b**), **Zin(2)** (**c**) or **Zin(3)** (**d**). Cells were incubated for 60 min in the presence and absence of Zn$^{2+}$ before CLSM imaging. Scale bar = 10 μm.

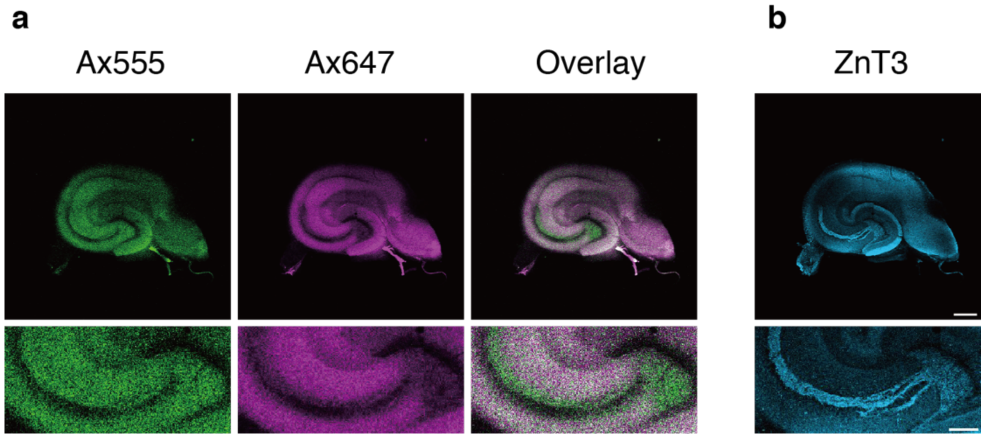


**Supplementary Fig. 4 | Visualization of activity-dependent $Zn^{2+}$ release at excitatory synapses in acutely prepared hippocampal slices. a**, CLSM imaging of the entire hippocampal slices labeled with **CAM2 (TCO, Ax555)** and **Zin(2)** after KCl depolarization. The slices were fixed with 4% PFA/PBS(–) containing 1 mM EDTA. **b**, CLSM imaging of a hippocampal slice immunostained with anti-ZnT3 (rabbit, Almone Labs AZT-013, ×1000). Scale bar = 500 µm (upper) and 200 µm (lower).

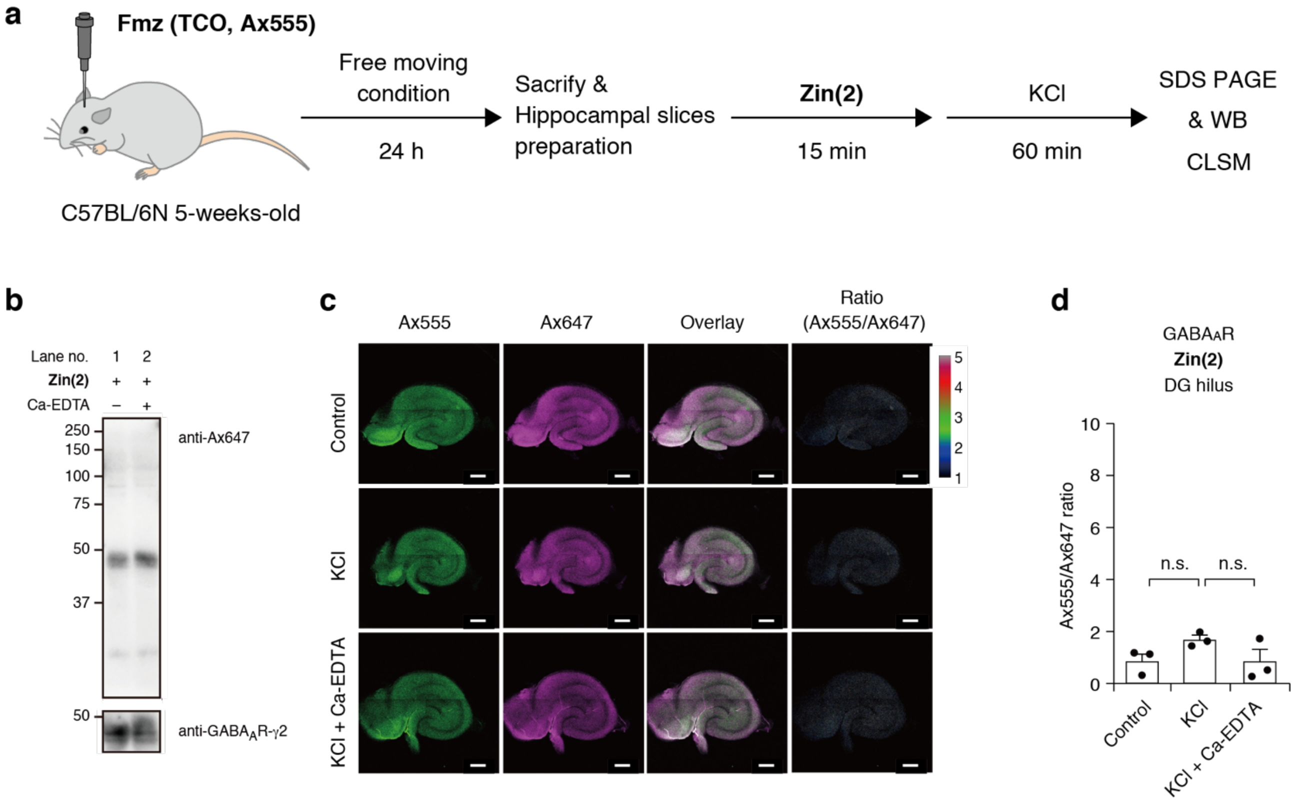


**Supplementary Fig. 5 | Evaluation of the local $Zn^{2+}$ release at GABAergic synapses in acute brain slices. a**, Experimental workflow in Supplementary Fig. 5. **Fmz (TCO, Ax555)** (100 μM, 4.5 μL) was directly injected into the lateral ventricle (LV) of a living mouse brain, and 24 h after injection, the hippocampus was isolated. Subsequently, acute hippocampal slices (300 μm in thickness) were prepared and incubated with **Zin(2)** (1 μM) for 15 min. After labeling reaction, the slices were treated with 50 mM KCl in the presence and absence of Ca-EDTA for 60 min. **b**, Western blot analysis of **Zin(2)** labeling to $GABA_A$Rs. After incubation with **Zin(2)**, the slices were washed and further incubated in ACSF for 60 min. In lane 2, 10 mM Ca-EDTA was co-incubated with **Zin(2)**. **c**, CLSM imaging of the hippocampal slices labeled with **Zin(2)**. The slices were fixed with 4% PFA/PBS(–) containing 1 mM EDTA. Scale bar = 500 μm. **d**, The Ax555/Ax647 ratio values in the hilus of DG region in Supplementary Fig. 5c. Data represents mean values ± s.e.m. (n = 3). Student's unpaired t-test with a two tailed distribution. n.s.: not significant ($p > 0.05$).

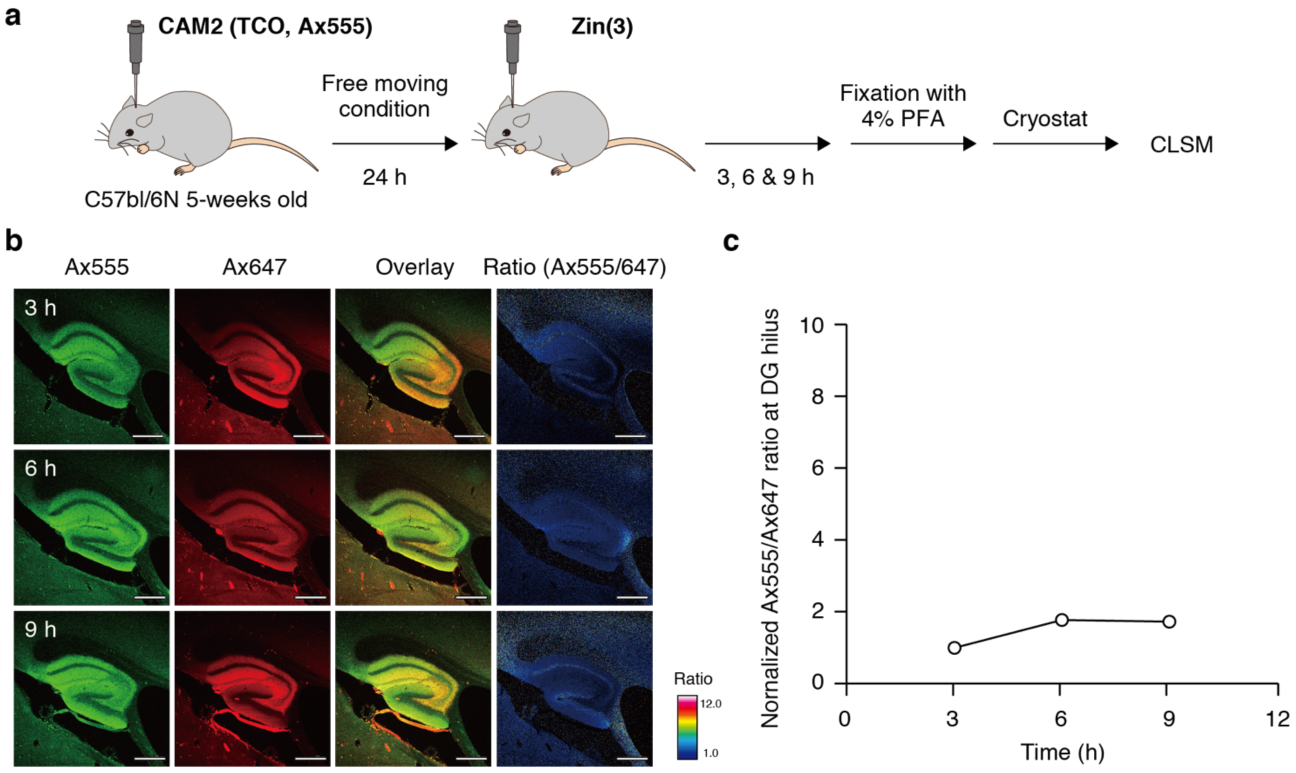


**Supplementary Fig. 6 | Evaluation of basal $Zn^{2+}$ concentrations at excitatory synapses in mouse hippocampus. a**, Experimental workflow in Supplementary Fig. 6. **CAM2 (TCO, Ax555)** (100 µM, 4.2 µL) was directly injected into the lateral ventricle (LV) of a living mouse brain, and 24 h after the injection, **Zin(3)** (100 µM, 4.2 µL) was injected into the LV. The mice were perfused transcardially with 4% PFA/PBS(–) containing 1 mM EDTA at 3, 6 and 9 h after the injection of **Zin(3)**. **b**, CLSM imaging of hippocampus area of sagittal cryosections of mouse brain samples labeled with **CAM2 (TCO, Ax555)** and **Zin(3)**. CLSM imaging was conducted using LSM800 (Zeiss) equipped with a 5× objective (NA = 0.25). scale bar = 500 µm. **c**, Plot of normalized Ax555/Ax647 ratio values at 3, 6 and 9 h after the injection of **Zin(3)**. The Ax555/Ax647 ratio value at 3 h after the **Zin(3)** injection was defined as 1.

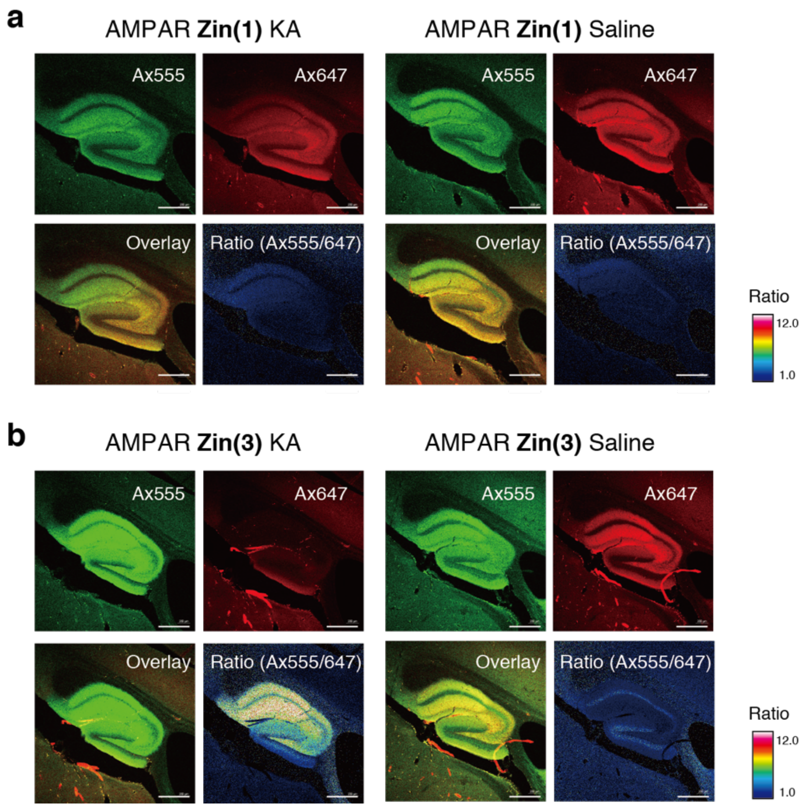


**Supplementary Fig. 7 | Recording of activity dependent synaptic $Zn^{2+}$ release in living mouse brains labeled with CAM2 (TCO, Ax555)/Zin(1) or CAM2 (TCO, Ax555)/Zin(3). a**, CLSM imaging of hippocampus region of mouse brain cryosections labeled with **CAM2 (TCO, Ax555)** and **Zin(1)**. The mice were injected intraperitoneally with kainate (KA) (25 mg/kg mouse body weight) or saline 3 h before the perfusion with cold 4% PFA/PBS(-) containing 1 mM EDTA. Scale bar = 500 μm. **b**, CLSM imaging of hippocampus region of mouse brain cryosections labeled with **CAM2 (TCO, Ax555)** and **Zin(3)**. The mice were injected intraperitoneally with KA (25 mg/kg mouse body weight) or saline 3 h before the perfusion with cold 4% PFA/PBS(–) containing 1 mM EDTA. Scale bar = 500 μm.

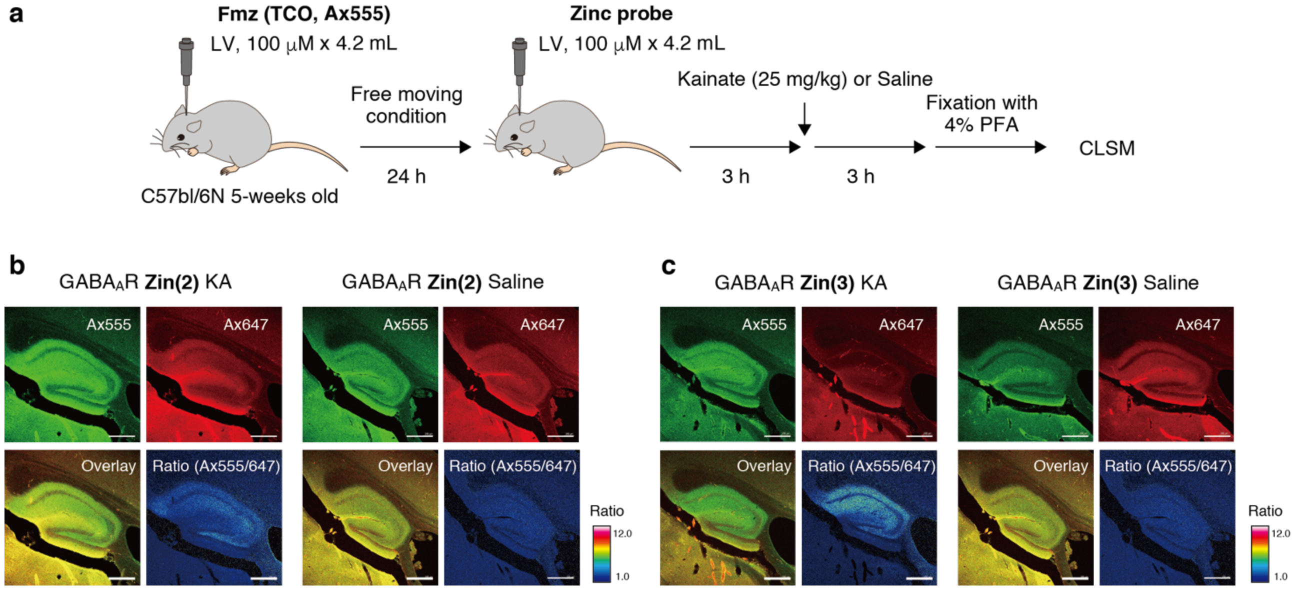


**Supplementary Fig. 8 | Evaluation of $Zn^{2+}$ leaking into GABAergic synapses in living mouse brains. a**, Experimental workflow in Supplementary Fig. 8. $GABA_AR$-based $Zn^{2+}$ sensors were constructed as described in Fig. 5. Three hours after the injection of **Zin(2)** or **Zin(3)** into lateral ventricle (LV) of a living mouse brain, the mice were injected intraperitoneally with kainate (KA) (25 mg/kg mouse body weight) or saline. The mice were perfused transcardially with 4% PFA/PBS(–) containing 1 mM EDTA at 3 h after the KA injection. CLSM imaging was conducted using a 5× (NA, 0.25) objective. **b**, CLSM imaging of hippocampus area of mouse brain cryosections labeled with **Fmz (TCO, Ax555)** and **Zin(2)**. The mice were injected intraperitoneally with KA or saline 3 h before the PFA perfusion. Scale bar = 500 μm. **c**, CLSM imaging of hippocampus region of mouse brain cryosections labeled with **Fmz (TCO, Ax555)** and **Zin(3)**. The mice were injected intraperitoneally with KA or saline 3 h before the PFA perfusion. Scale bar = 500 μm.

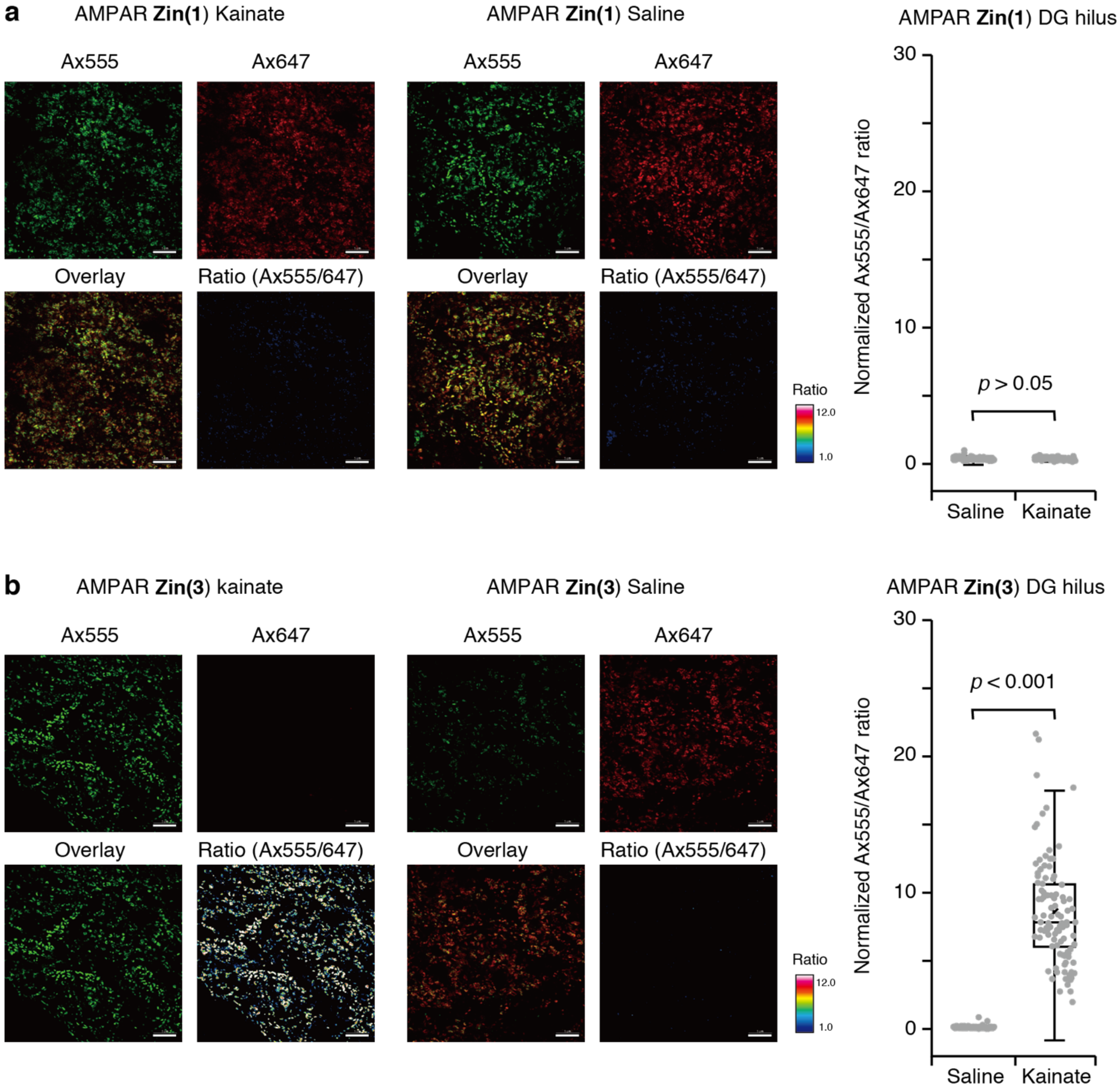


**Supplementary Fig. 9 | High-resolution images of the hippocampal DG hilus region of mouse brain cryosections. a**, CLSM images of the hilus area of hippocampal dentate gyrus (DG) labeled with **CAM2 (TCO, Ax555)** and **Zin(1)**. The mice were injected intraperitoneally with kainate (KA) (25 mg/kg mouse body weight) or saline 3 h before the perfusion with cold 4% PFA/PBS(–) containing 1 mM EDTA. Scale bar = 5 µm. Boxplot of the Ax555/Ax647 ratio values of the fluorescence bright spots. The number of ROIs is 100 for each of the images (n = 100). The horizonal line and × within each box indicate the median and mean, respectively. Boxes show interquartile range (IQR). Whiskers show 1.5 × IQR. Student's unpaired t-test with a two tailed distribution. **b**, CLSM images of the hilus area of hippocampal DG labeled with **CAM2 (TCO, Ax555)** and **Zin(3)**. The mice were injected intraperitoneally with KA (25 mg/kg mouse body weight) or saline 3 h before the perfusion with cold 4% PFA/PBS(–) containing 1 mM EDTA. scale bar = 5 μm. Boxplot of the Ax555/Ax647 ratio values of the fluorescence bright spots. The number of ROIs is 100 for each of the images (n = 100). The horizonal line and × within each box indicate the median and mean, respectively. Boxes show interquartile range (IQR). Whiskers show 1.5 × IQR. Student's unpaired t-test with a two tailed distribution.

**General materials and methods for organic synthesis**

All chemical reagents and solvents were obtained from commercial suppliers (Tokyo Chemical Industry (TCI), Sigma-Aldrich, Fujifilm, Watanabe Chemical Industries, Thermo Fisher Scientific) and used without further purification. Thin layer chromatography (TLC) was performed on silica gel 60 F254 precoated aluminum sheets (Merck) and visualized by fluorescence quenching, fluorescence upon 365 nm excitation, and TLC stains. Chromatographic purification was accomplished using a Biotage Isolera Spektra One Instrument installed with Sfär Silica HC D columns. $^{1}$H-NMR and $^{13}$C-NMR spectra were recorded in deuterated solvents on a Varian Mercury 400 (400 MHz) or a JEOL ECS400 (400 MHz) spectrometer. Chemical shifts were referenced to tetramethylsilane (0 ppm) or residual solvent peak. Multiplicities are abbreviated as follows: s = singlet, d = doublet, t = triplet, dd = doublet of doublets, td = triplet of doublets, m = multiplet. High-resolution mass spectra were measured on an Exactive (Thermo Scientific) equipped with electron spray ionization (ESI). Reversed-phase HPLC (RP-HPLC) was carried out on a Hitachi Chromaster system equipped with a 5410 UV detector (set at 220 nm) or a 5430 diode array detector and a YMC-Pack ODS-A column (20 × 250 mm) or a COSMOSIL 5C18-AR-II column (10 × 250 mm).

**Abbreviation**

Ax647: Alexa Fluor 647

Ax555: Alexa Fluor 555

Boc: tert-butoxycarbonyl

DCM: dichloromethane

DIPEA: *N*,*N*-diisopropylethylamine

DMF: *N*,*N*-dimethylformamide

DMSO: dimethyl sulfoxide

EDC: 1-ethyl-3-(3-dimethylaminopropyl)carbodiimide

EtOAc: ethyl acetate

HBTU: 2-(1H-benzotriazol-1-yl)-1,1,3,3-tetramethyluronium hexafluorophosphate

HOBt: 1-hydroxybenzotriaole

MeTz: methyltetrazine

NHS: *N*-hydroxysuccinimide

TCO: *trans*-cyclooctene

TEA: triethylamine

TFA: trifluoroacetic acid

TSTU: *N,N,N',N'*-tetramethyl-*O*-(*N*-succinimidyl)uranium tetrafluotoborate

TEAA: triethylammonium acetate

MeTz-$NH_2$: methyltetrazine amine

**Synthesis of 1**

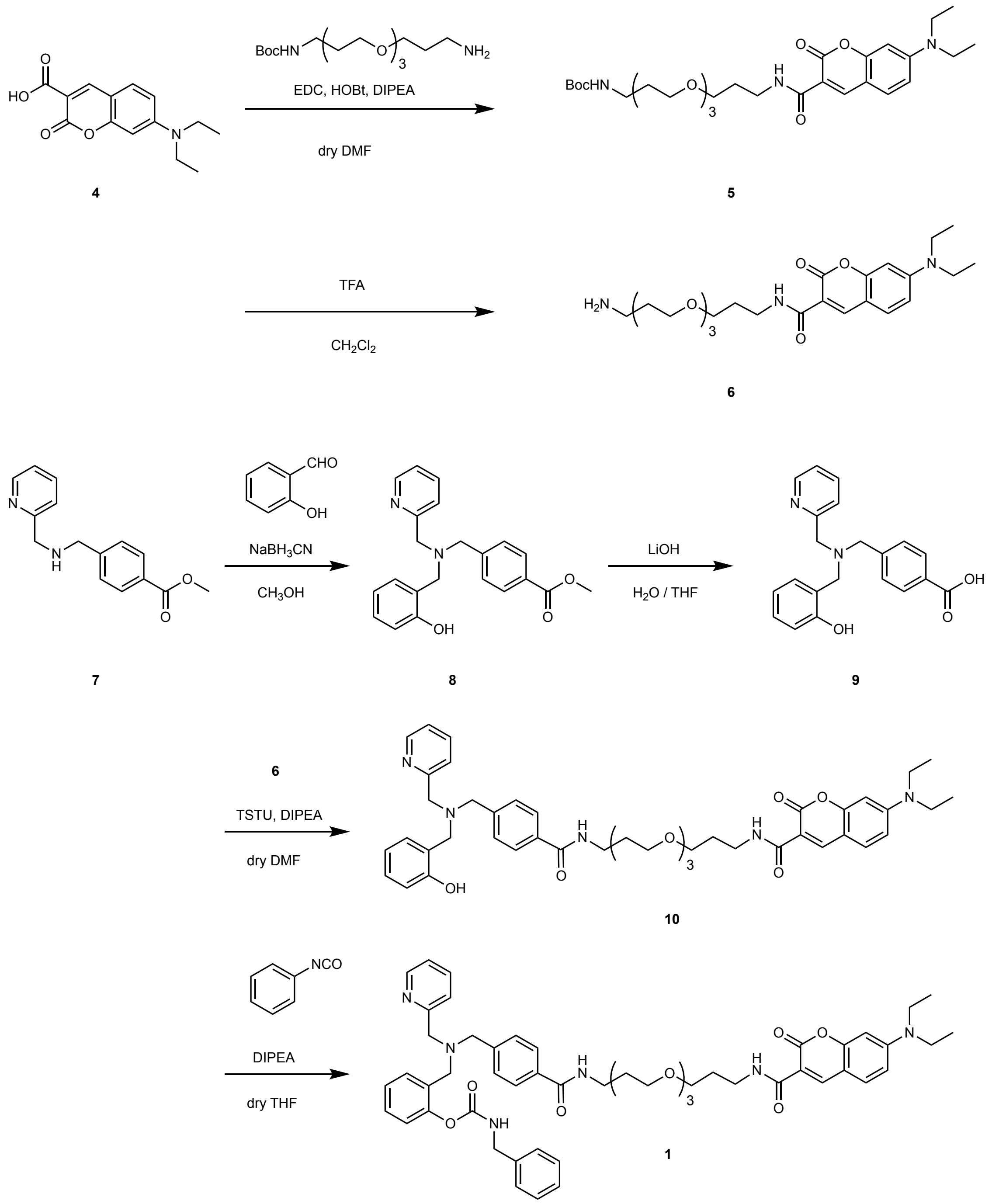


**5:** To a solution of **4** (70 mg, 268 μmol, 1.0 eq) in dry DMF, Boc-dPEG3-$NH_2$ (103 mg, 322 μmol, 1.2 eq), HOBt (47 mg, 348 μmol, 1.3 eq), EDC HCl (66.7 mg, 348 μmol, 1.3 eq) and DIPEA (186 μL, 1.07 mmol, 4.0 eq) were added. After stirring at r.t. under $N_2$ overnight, the reaction mixture was diluted with 20 mL of $CH_2Cl_2$ and washed with 5% citric acid aq. (20 mL×1), sat. $NaHCO_3$ aq. (20 mL×1), $H_2O$ (20 mL×1), and brine (20 mL×1). The organic layer was dried over $Na_2SO_4$. After evaporating the solvent, **5** (153.6 mg, 272 μmol, 101%) was obrained as yellow oil. $^1$H NMR (400

MHz, $CDCl_3$): $\delta$ 8.90 (t, $J$ = 3.6 Hz, 1H), 8.70 (s, 1H), 7.43 (d, $J$ = 6.0 Hz, 1H), 6.65 (dd, $J$ = 6.0, 1.6 Hz, 1H), 6.50 (d, $J$ = 1.6 Hz, 1H), 3.70 (m, 2H), 3.67 (m, 2H), 3.64 (m, 2H), 3.60 (m, 4H), 3.56 (m, 4H), 3.47 (q, $J$ = 4.8 Hz, 4H), 3.25 (q, $J$ = 4.0 Hz, 2H), 1.93 (m, 2H), 1.43 (s, 9H), 1.25 (t, $J$ = 4.8 Hz, 6H). $^{13}C$ NMR (150 MHz, $CDCl_3$): $\delta$ 163.1, 162.7, 157.6, 156.1, 152.4, 148.0, 131.1, 110.4, 109.9, 108.4, 96.5, 78.8, 70.6, 70.6, 70.4, 70.2, 69.7, 69.2, 45.1, 38.6, 37.1, 29.6, 29.5, 28.5, 12.4.

**6:** To a solution of compound **5** in 2 mL of $CH_2Cl_2$, 0.5 mL of TFA was added on ice. After stirring for 2 h, the solution was azoetroped with toluene (×3). **6** (TFA salt, quant) was obtaind as yellow oil. The product was directly used for the next reactions without further purification.

**8:** To a solution of compound **7**[S1] (120 mg, 468 µmol, 1.0 eq) in 4 mL of $CH_3OH$, salicylaldehyde (114 mg, 0.936 mmol, 2.0 eq) and $NaBH_3CN$ (198 mg, 0.936 mmol, 2.0 eq) were added. After adjusting the solution pH to 6 with AcOH (5-6 drops), the reaction was stirred at r.t. overnight. The unreacted $NaBH_3CN$ was quenched with 1 M HCl aq. and the solution was allowed to stir for another 1 h. After filtration, the solvent was evaporated and the crude product was purified by flash column chromatography on silica gel (0–10 % $CH_3OH/CHCl_3$). **8** (149.3 mg, 412 µmol, 88.0%) was obtained as pale yellow oil. $^1H$ NMR (400 MHz, $CDCl_3$): $\delta$ 8.61 (dd, $J$ = 4.4, 1.6 Hz, 1H), 7.97 (d, $J$ = 8.4 Hz, 2H), 7.66 (td, $J$ = 7.6, 1.6 Hz, 1H), 7.40 (d, $J$ = 8.0 Hz, 2H), 7.21 (m, 3H), 7.03 (d, $J$ = 7.6Hz, 1H), 6.90 (d, $J$ = 8.4 Hz, 1H), 6.79 (t, $J$ = 7.6 Hz, 1H), 3.87 (s, 3H), 3.76 (s, 2H), 3.75 (s, 2H), 3.71 (s, 2H). $^{13}C$ NMR (150 MHz, $CDCl_3$): $\delta$ 166.9, 157.5, 157.4, 149.1, 143.1, 136.8, 129.8, 129.8, 129.2, 129.1, 123.2, 122.5, 122.3, 119.0, 116.5, 58.6, 57.6, 57.1, 52.1.

**9:** To a solution of compound **8** (21.8 mg, 62.6 µmol, 1.0 eq) in 600 µL of $THF/H_2O$ (1/1, v/v), 1 M LiOH aq. (188 µL, 188 µmol, 3.0 eq) was added on ice. The solution was stirred at r.t. for 2.5 h. After diluting with 20 mL of $H_2O$, the solution pH was adjusted to 6 with 1 M HCl aq. The product was extracted with chloroform (20 mL×5) and the organic layer was dried over $Na_2SO_4$. After evaporation, **9** (20.5 mg, 58.8 µmol, 94.0%) was obtained as white oily solids. The product was directly used for the next reaction without further purification.

**10:** To a solution of **9** (10 mg, 28.7 µmol, 1.0 eq) in 0.3 mL of dry DMF, TSTU (13 mg, 43.1 µmol, 1.5 eq) and DIPEA (50 µL, 287 µmol, 10 eq) were added. After stirring at r.t. under $N_2$ for 5 min, **6**

(26.6 mg, 47.3 μmol, 1.6 eq) was added and stirred at r.t. under $N_2$ overnight. The solution was diluted with 20 mL of $CHCl_3$ and washed with 5% citric acid aq. (20 mL×1), sat. $NaHCO_3$ aq. (20 mL×1), and brine (20 mL×1). The organic layer was dried over $Na_2SO_4$. The crude product was purified by flash column chromatography on silica gel (0-10 % $CH_3OH/CHCl_3$). **10** (with impurities, 8.1 mg, 10.2 μmol, 35.5%) was obtained as yellow oil. $^1$H-NMR ($CDCl_3$, 500 MHz): $\delta$ 8.68 (s, 1H), 8.61-8.59 (m, 1H), 7.75 (d, $J$ = 4.8 Hz, 2H), 7.64 (t, $J$ = 6.4 Hz, 1H), 7.46-7.37 (m, 2H), 7.28-7.15 (m, 3H), 7.00 (t, $J$ = 6.4 Hz, 1H), 6.87 (d, $J$ =6.4 Hz, 1H), 6.76 (t, $J$ = 6.8 Hz, 2H), 6.62 (m, 1H), 6.49 (d, $J$ =2.0 Hz, 1H), 3.75-3.35 (m, 26H), 2.14-1.77 (m, 4H), 1.31-1.20 (m, 6H).

**1:** To a solution of **10** (8.1 mg, 10.2 μmol, 1.0 eq) in 0.8 mL of dry THF, benzyl isocyanate (1.4 mg, 10.2 μmol, 4.0 eq) and DIPEA (5.3 μL, 30.6 μmol, 5.0 eq) were added. After stirring under $N_2$ at r.t. overnight, the crude product was purified by RP-HPLC (A/B = 30/70 to 66/34 over 54 min, A: $CH_3CN$ (0.1% TFA), B: $H_2O$ (0.1% TFA)). After lyophilization, **1** (1.02 μmol, 10.0%) was obtained as yellow solid. The amount of **1** was determined by UV-Vis ($\varepsilon$ = 49000 at 419 nm in $CH_3OH$). HR-ESI MS *m/z* calcd. for $C_{53}H_{62}N_6O_9Na$ $[M + Na]^+$ 949.4470, found 949.4467. The purity of **1** was checked by HPLC as below ($CH_3CN$ (0.1% TFA)/$H_2O$ (0.1% TFA) = 30/70 (0 min) → 70/30 (30 min)). $^1$H-NMR (400 MHz, $CDCl_3$): $\delta$ 8.62 (s, 1H), 8.50 (m, 1H), 7.77 (d, $J$ = 8.0 Hz, 2H), 7.62-7.00 (m, 15H), 6.62 (d, 1H), 6.48 (s, 1H), 4.58 (s, 2H), 4.40 (s, 2H), 4.30 (d, 2H), 3.96 (s, 2H), 3.66-3.42 (m, 20H), 1.22 (m, 6H).

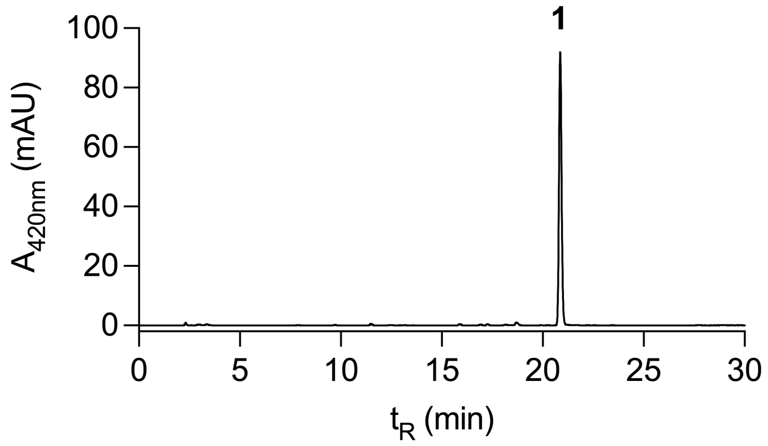

**Synthesis of 2**

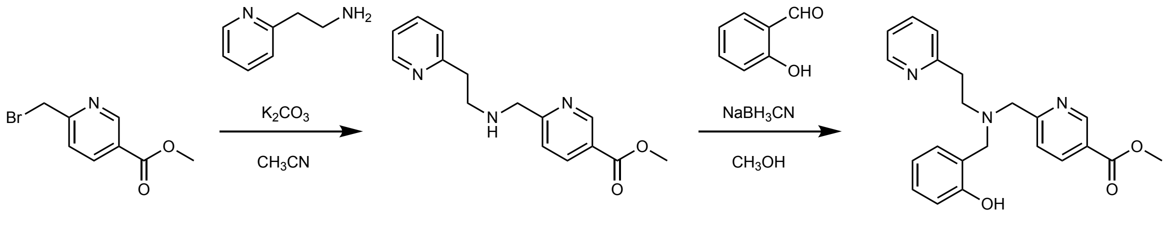


11 12 13

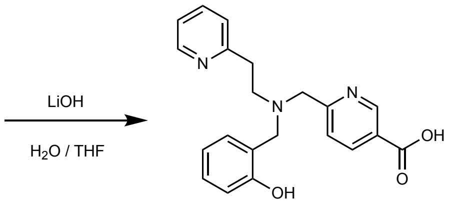


14

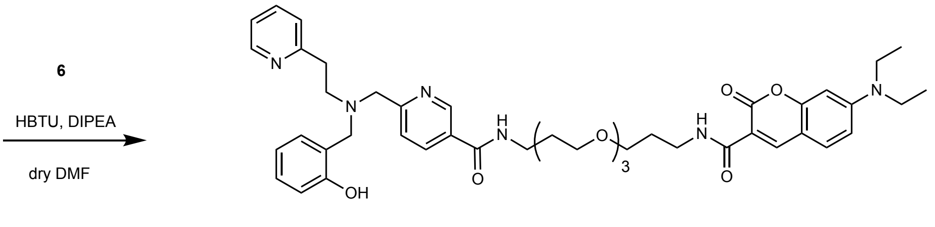


15

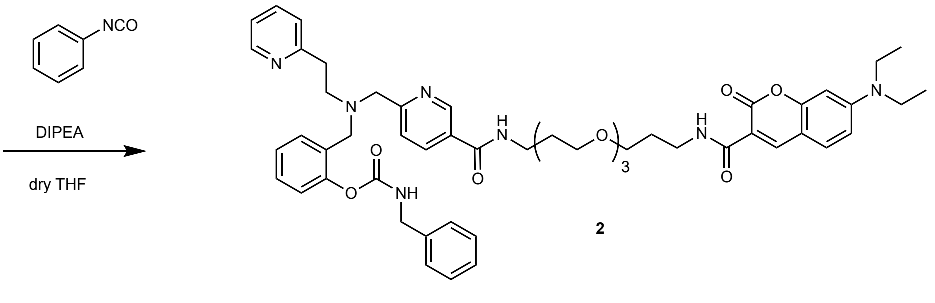


2

**12:** To a solution of 2-(2-aminoethyl)pyridine (777.8 μL, 6.5 mmol, 5.0 eq) and $K_2CO_3$ (215.6 mg, 1.6 mmol, 1.2 eq) in $CH_3CN$ (15 mL), **11**[S2,S3] (299.1 mg, 1.3 mmol, 1.0 eq) in $CH_3CN$ (20 mL) was added dropwise on ice. After stirring at r.t. overnight, the crude product was purified by flash column chromatography on silica gel (0–10 % $CH_3OH/CHCl_3$) to yield **12** (275.2 mg, 1.01 mmol, 78.0 %) as a yellow oil. $^{1}$H NMR (400 MHz, $CDCl_3$): $\delta$ 9.12 (d, $J$ = 1.6 Hz, 1H,), 8.54 (d, $J$ = 8.4 Hz, 1H), 8.23 (dd, $J$ = 8.4, 2.4 Hz, 1H), 7.62 (td, $J$ = 7.6, 2.0 Hz, 1H), 7.40 (d, $J$ = 8.0 Hz, 1H), 7.18 (d, $J$ = 8.0 Hz, 1H), 7.14 (td, $J$ = 6.4, 1.2 Hz, 1H), 4.01 (s, 2H), 3.93 (s, 3H), 3.12 (d, $J$ = 7.6 Hz, 2H), 2.93 (d, $J$ = 6.4 Hz, 2H). $^{13}$C NMR (150 MHz, $CDCl_3$): $\delta$ 176.0, 165.5, 158.7, 150.5, 148.9, 138.1, 137.1, 125.2, 123.6, 122.5, 121.9, 52.5, 50.7, 47.5, 35.0.

**13:** To a solution of **12** (275.2 mg, 1.0 mmol, 1.0 eq) in $CH_3OH$ (10 mL), 2-hydroxybenzaldehyde (127.1 μL, 1.2 mmol, 1.2 eq) and $NaBH(OAc)_3$ (215.0 mg, 1.0 mmol, 1.0 eq) were added, the mixture was stirred at r.t. overnight. The unreacted $NaBH(OAc)_3$ was quenched with 1 M HCl aq. and the solution was allowed to stir for another 1 h. The solvent was evaporated and the crude product was purified by flash column chromatography on silica gel (0–15 % $CH_3OH/CHCl_3$) to yield compound **13** (297.0 mg, 0.78 mmol, 78.0 %) as a yellow oil. $^1$H NMR (400 MHz, $CDCl_3$): $\delta$ 9.14 (d, $J$ = 1.6 Hz, 1H), 8.49 (d, $J$ = 4.0 Hz, 1H), 8.18 (dd, $J$ = 8.4, 2.4 Hz, 1H), 7.60 (td, $J$ = 7.6, 1.6 Hz, 1H), 7.22 (d, $J$ = 8.4 Hz, 1H), 7.19-7.12 (m, 2H), 7.07 (d, $J$ = 7.6 Hz, 1H), 7.03 (dd, $J$ = 7.2, 1.6 Hz, 1H), 6.83 (dd, $J$ = 8.0, 0.8 Hz, 1H), 6.80 (td, $J$ = 7.6, 1.2 Hz, 1H), 3.94 (s, 3H), 3.90 (s, 2H), 3.87 (s, 2H), 3.05 (d, $J$ = 6.4 Hz, 2H), 3.02 (d, $J$ = 6.8 Hz, 2H). $^{13}$C NMR (150 MHz, $CDCl_3$): $\delta$ 165.6, 162.3, 159.2, 157.4, 150.3, 149.0, 137.7, 136.8, 129.4, 129.0, 124.7, 123.5, 122.8, 122.1, 121.6, 119.2, 116.4, 59.2, 57.6, 53.8, 52.4, 35.0.

**14:** To a solution of **13** (121.9 mg, 323 μmol, 1.0 eq) in 6 mL of $THF/H_2O$ (1/1, v/v) on ice, 1 M LiOH aq. (969 μL, 969 μmol, 3.0 eq) was added. The solution was stirred at r.t. for 2.5 h. After diluting with 20 mL of $H_2O$, the solution pH was adjusted to 6 with 1 M HCl aq. The product was extracted with $CHCl_3$ (20 mL×5) and the organic layer was dried over $Na_2SO_4$. After evaporation, **14** (94.8 mg, 261 μmol, 80.8%) was obtained as colorless oil. The product was directly used for the next reaction without further purification.

**15:** To a solution of **14** (10 mg, 27.5 μmol, 1.1 eq) in 0.8 mL of dry DMF, **6** (14 mg, 24.9 μmol, 1.0 eq), HBTU (16 mg, 41 μmol, 1.6 eq) and DIPEA (48 μL, 275 μmol, 11 eq) were added. The reaction mixture was allowed to stir under $N_2$ overnight. After evaporating the solvent, the crude product was roughly purified by flash column chromatography on silica gel ($CH_3OH/CHCl_3$) and used for the next reaction. $^1$H-NMR (400 MHz, $CDCl_3$): $\delta$ 8.96 (s, 1H), 8.69 (s, 1H), 8.65 (s, 1H), 8.47 (d, $J$ = 4.8 Hz, 1H), 8.06 (d, $J$ = 8.4 Hz, 1H), 7.56 (t, $J$ = 8.0 Hz, 1H), 7.51 (s, 1H (amide)), 7.44 (m, 1H), 7.39 (dd, $J$ = 8.8, 1.6 Hz, 1H), 7.19-7.12 (m, 2H), 7.05 (d, $J$ = 8.0 Hz, 1H), 7.00 (d, $J$ = 7.2 Hz, 1H), 6.79 (t, $J$ = 8.8 Hz, 1H), 6.64 (t, $J$ = 9.2 Hz, 1H), 6.48 (d, $J$ = 10.0 Hz, 1H), 3.88 (s, 2H), 3.85 (s, 2H), 3.68-3.36 (20H, m), 3.03-2.99 (m, 4H), 1.85-1.77 (m, 4H), 1.24 (m, 6H).

**2:** To a solution of **15** (crude product) in 0.8 mL of dry THF, benzyl isocyanate (1.4 mg, 10.2 μmol, 4.0 eq) and DIPEA (5.3 μL, 30.6 μmol, 5.0 eq) were added. The reaction mixture was allowed to stir under $N_2$ at r.t. overnight. The crude product was purified by RP-HPLC (A/B = 30/70 to 70/30 over 40 min, A: $CH_3CN$ (0.1% TFA), B: $H_2O$ (0.1% TFA)). After lyophilization, **2** (0.75 μmol, 2.7% in 2 steps) was obtained as yellow solid. The amount of **2** was determined by UV-Vis ($\varepsilon$ = 49000 at 419 nm in $CH_3OH$). HR-ESI MS *m/z* calcd. for $C_{53}H_{64}N_7O_9$ $[M+H]^+$ 942.4760, found 942.4735. The purity of **2** was checked by HPLC as below ($CH_3CN$ (0.1% TFA)/$H_2O$ (0.1% TFA) = 30/70 (0 min) → 70/30 (30 min)). $^1$H-NMR (400 MHz, $CDCl_3$): $\delta$ 8.89 (s, 1H), 8.67 (1H, s), 8.62 (1H, s), 8.14 (m, 1H), 8.00 (m, 1H), 7.51-6.95 (m, 16H), 6.61 (td, $J$ = 10.0, 2.4 Hz, 1H), 6.46 (dd, $J$ = 10.0, 1.6 Hz 1H), 4.38 (d, $J$ = 6.0 Hz, 2H), 3.80 (s, 2H), 3.73 (s, 2H), 3.69-3.34 (m, 20H), 3.04 (t, $J$ = 7.2 Hz, 2H), 2.79 (m, 2H), 1.82-1.74 (m, 4H), 1.22 (t, $J$ = 6.8 Hz, 6H).

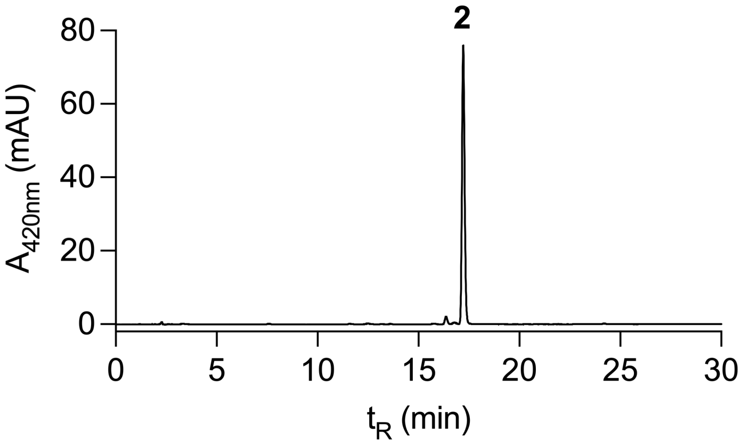

**Synthesis of 3**

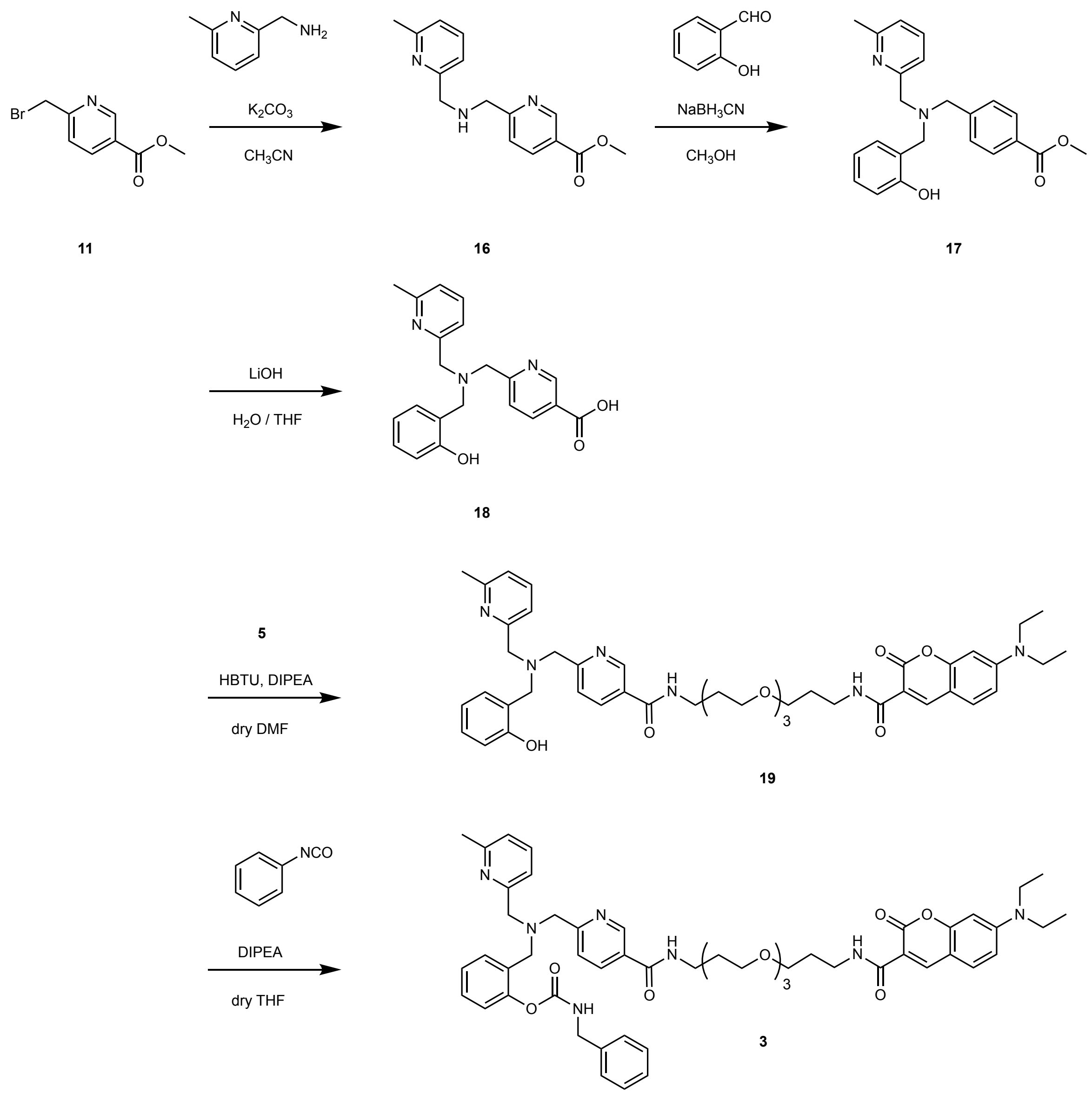


**16:** To a solution of 6-methylpyridin-2-yl-methanamine (250 mg, 2.0 mmol, 5.0 eq) and $K_2CO_3$ (52.1 mg, 0.377 mmol, 0.9 eq) in $CH_3CN$ (10 mL), **11** (91.7 mg, 0.4 mmol, 1.0 eq) in $CH_3CN$ (15 mL) was added dropwise on ice. After stirring at r.t. overnight, the resulting precipitate was filtered through celite and the solvent was evaporated. The crude product was purified by flash column chromatography on silica gel (0–10 % $CH_3OH/CHCl_3$) to yield **16** (119.3 mg, 440 μmol, 110%) as a brown oil. $^1H$ NMR (400 MHz, $CDCl_3$): $\delta$ 9.15 (d, $J$ = 2.0 Hz, 1H), 8.25 (d, $J$ = 8.0 Hz, 1H), 7.53 (t, $J$ = 7.6 Hz, 1H), 7.47 (d, $J$ =8.0 Hz, 1H), 7.13 (d, $J$ = 7.6 Hz, 1H), 7.03 (d, $J$ = 7.6 Hz, 1H), 4.08 (s, 2H), 3.97 (s, 2H), 3.94 (s, 3H), 2.54 (s, 3H).

**17:** To a solution of **16** (120 mg, 453 μmol, 1.0 eq) in $CH_3OH$ (4 mL), 2-hydroxybenzaldehyde (111 mg, 906 μmol, 2.0 eq) and $NaBH_3CN$ (57 mg, 906 μmol, 2.0 eq) were added. The solution pH was adjusted to 6 with AcOH, and the reaction mixture was stirred at r.t. overnight. The unreacted $NaBH_3CN$ was quenched with 1M HCl aq. and the solution was allowed to stir for another 1 h. After filtration, the solvent was evaporated and the crude product was purified by flash column chromatography on silica gel (0–10 % $CH_3OH/CHCl_3$) to yield **17** (97.3 mg, 258 μmol, 57.0%) as a brown oil. $^1$H NMR (400 MHz, $CDCl_3$): $\delta$ 9.13 (d, $J$ = 2.0 Hz, 1H), 8.21 (dd, $J$ = 8.0, 2.0 Hz, 1H), 7.53-7.49 (m, 2H), 7.17 (td, $J$ = 7.6, 1.6 Hz, 1H), 7.08 (d, $J$ = 7.6Hz, 1H), 7.06-7.01 (m, 2H), 6.91 (dd, $J$ = 8.0, 1.2 Hz, 1H), 6.77 (td, $J$ = 7.6, 1.2 Hz, 1H), 3.92-3.90 (m, 5H), 3.82 (s, 2H), 3.77 (s, 2H), 2.59 (s, 3H).

**18:** To a solution of **17** (97.3 mg, 258 μmol, 1.0 eq) in 6 mL of THF/$H_2O$ (1/1, v/v) on ice, 1 M LiOH aq. (774 μL, 774 μmol, 3.0 eq) was added. The solution was stirred at r.t. overnight. After diluting with 20 mL of $H_2O$, the solution pH was adjusted to 6 with 1 M HCl aq. The product was extracted with $CHCl_3$ (20 mL×5). The organic layer was dried over $Na_2SO_4$. After evaporation, the compound **18** (76.1 mg, 209 μmol, 81.0%) was obtained as colorless oil. The product was directly used for the next reaction without further purification.

**19:** To a solution of **18** (13.7 mg, 37.7 μmol, 1.0 eq) in 0.5 mL of dry DMF, **7** (26.2 mg, 56.6 μmol, 1.5 eq), HBTU (18.6 mg, 48.0 μmol, 1.3 eq) and DIPEA (32.9 μL, 189 μmol, 5.0 eq) were added. The reaction mixture was allowed to stir under Ar overnight. After evaporating the solvent, the crude product was redissolved in 30 mL of EtOAc and washed with 5% citric acid aq. (25 mL×2), sat. $NaHCO_3$ aq. (25 mL×2), water (25 mL×1) and brine (25 mL×1). The organic layer was dried over $Na_2SO_4$. After evaporation, **19** (with impurities, 12.6 mg, 15.6 μmol, 41.4%) was obrained as yellow oil. $^1$H-NMR ($CDCl_3$, 400MHz): $\delta$ 8.97 (d, $J$ =1.6 Hz, 1H), 8.87 (t, $J$ = 6.0 Hz, 1H (amide)), 8.64 (s, 1H), 8.09 (dd, $J$ = 8.0, 2.4 Hz, 1H), 7.52-7.37 (m, 4H), 7.16 (td, $J$ = 7.6, 1.6 Hz, 1H), 7.08 (d, $J$ =7.6 Hz, 1H), 7.05-7.00 (m, 2H), 6.88 (dd, $J$ = 8.0, 1.2 Hz, 1H), 6.75 (td, $J$ = 7.2, 1.2 Hz, 1H), 6.62 (dd, $J$ = 8.8, 2.0 Hz, 1H), 6.48 (dd, $J$ = 9.2, 2.4 Hz, 1H), 3.88 (s, 2H), 3.81 (s, 2H), 3.76 (s, 2H), 3.68-3.41 (m, 20H), 2.58 (s, 3H), 1.90-1.80 (m, 4H), 1.25-1.21 (m, 6H).

**3:** To a solution of **19** (12.6 mg, 15.6 μmol, 1.0 eq) in 1 mL of dry THF, benzyl isocyanate (8.3 mg,

62.4 μmol, 4.0 eq) and DIPEA (11 μL, 62.4 μmol, 4.0 eq) were added. The reaction mixture was allowed to stir under Ar at r.t. overnight. The crude product was purified by RP-HPLC (A/B = 30/70 to 70/30 over 40 min, A: $CH_3CN$ (0.1% TFA), B: $H_2O$ (0.1% TFA)). After lyophilization, **3** (0.44 μmol, 2.8%) was obtained as yellow solid. The amount of **3** was determined by UV-Vis ($\varepsilon$ = 49000 at 419 nm in $CH_3OH$). HR-ESI MS *m/z* calcd. for $C_{53}H_{64}N_7O_9$ $[M+H]^+$ 942.4760, found 942.4757. The purity of **3** was checked by HPLC as below ($CH_3CN$ (0.1% TFA)/$H_2O$ (0.1% TFA) = 0/100 (0 min) → 10/90 (5 min) → 50/50 (30 min)). $^1$H-NMR ($CDCl_3$, 400MHz): $\delta$ 9.12 (s, 1), 8.90 (t, $J$ = 5.6 Hz, 1H (amide)), 8.61 (s, 1H), 8.32 (dd, $J$ = 8.0, 2.0 Hz, 1H), 7.99 (t, $J$ = 5.2 Hz, 1H (amide)), 7.75-7.71 (m, 2H), 7.57-7.52 (m, 2H), 7.39-6.97 (m, 10H), 6.63 (dd, $J$ = 8.8, 2.4 Hz, 1H), 6.46 (d, $J$ = 2.0Hz, 1H), 4.41 (d, $J$ = 4.8 Hz, 2H), 4.31 (s, 2H), 4.24 (s, 2H), 3.92 (s, 2H), 3.66-3.43 (m, 20H), 2.68 (s, 3H), 1.91-1.82 (m, 4H), 1.25-1.23 (m, 6H).

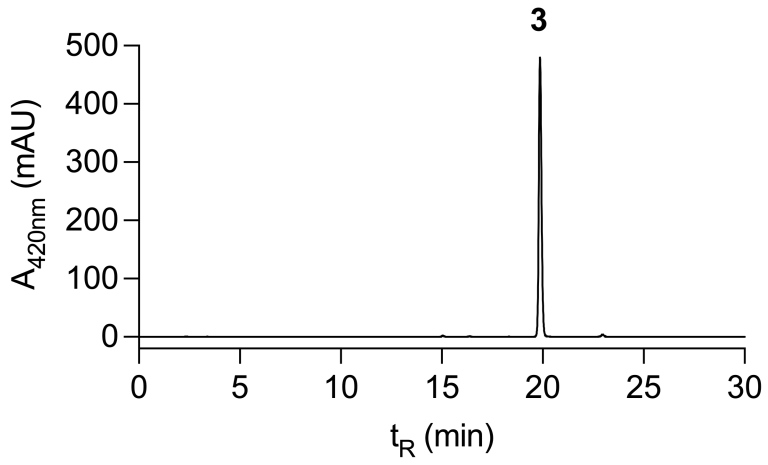

#### Synthesis of Zin(1)

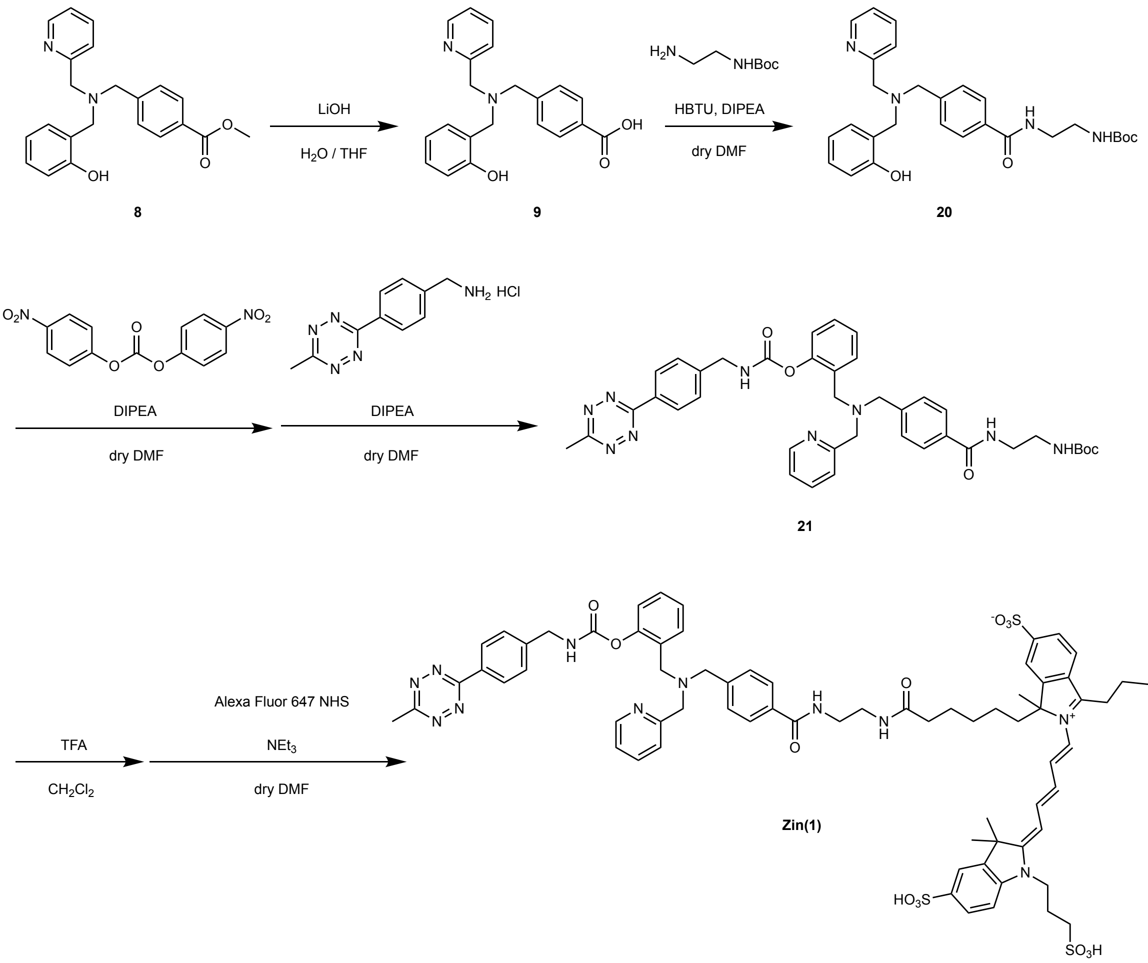


**20:** To a solution of **8** (135 mg, 0.37 mmol, 1.0 eq) in 10 mL of $H_2O/CH_3OH$ (1/4, v/v), $LiOH \cdot H_2O$ (93.6 mg, 2.24 mmol, 6.0 eq) was added. After stirring at r.t. for 20 h, the reaction solution was neutralized with 2 M HCl aq. and concentrated *in vacuo.* The residue was purified by flash column chromatography on silica gel (EtOAc) to give **9** (83.4 mg). To a solution of **9** (83.4 mg, 0.24 mmol, 1.0 eq) in dry DMF (2 mL), HBTU (136 mg, 0.36 mmol, 1.5 eq) and DIPEA (125 µL, 0.72 mmol, 3.0 eq) were added. After 10 min, *N*-Boc-ethylenediamine (56.9 µL, 0.36 mmol, 1.5 eq) was added. The mixture was allowed to stir at r.t. for 16 h. After evaporation of the solvent, the residue was purified by flash column chromatography on silica gel (0–100 % EtOAc/hexane) and reprecipitation with $CH_2Cl_2$ to give **20** (135 mg, 0.28 mmol, 75.7 % in 2 steps) as yellow oil. $^1$H NMR (400 MHz, $CDCl_3$): $\delta$ 8.57 (d, $J$ = 2.8 Hz, 1H), 7.75 (d, $J$ = 4.2 Hz, 2H), 7.70 (td, $J$ = 4.2, 1.2 Hz, 1H), 7.40 (m, 3H), 7.25 (d, $J$ = 4.2 Hz, 1H), 7.19 (td, $J$ = 5.6, 1.2 Hz, 1H), 7.09 (d, $J$ = 4.8 Hz, 1H), 6.90 (d, $J$ = 5.2 Hz, 1H), 6.81 (td, $J$ = 4.8, 0.8 Hz, 1H), 3.93 (s, 2H), 3.89 (s, 2H), 3.85 (s, 2H), 3.53 (m, 2H), 3.38 (m, 2H), 1.41 (s, 9H). $^{13}$C NMR (150 MHz, $CDCl_3$): $\delta$ 167.9, 157.6, 156.9, 148.9, 137.4, 133.5, 130.2,

129.7, 128.4, 127.5, 126.2, 126.1, 123.4, 123.0, 119.5, 117.3, 111.2, 80.2, 57.9, 56.7, 55.5, 41.8, 39.8, 28.3.

**21:** To a solution of **20** (5.0 mg, 10.2 µmol, 1.0 eq) in dry DMF (100 µL), bis(4-nitrophenyl) carbonate (4.7 mg, 15.3 µmol, 1.5 eq) and DIPEA (5.2 µL, 30.6 µmol, 3.0 eq) were added. After stirring at r.t. for 2 h, MeTz-amine HCl salt (3.6 mg, 15.3 µmol, 1.5 eq) and DIPEA (2.6 µL, 15.3 µmol, 1.5 eq) were added. The reaction mixture was allowed to stir at r.t. overnight. After evaporating the solvent, the crude product was purified by flash column chromatography on silica gel (0–10 % $CH_3OH/CHCl_3$) to yield **21** (2.9 mg, 4 µmol, 39.6 %) as a pink solid. $^1$H NMR (400 MHz, $CDCl_3$): $\delta$ 8.58 (d, $J$ = 6.8 Hz, 2H), 8.43 (d, $J$ = 6.0 Hz, 1H), 7.75 (d, $J$ = 6.4 Hz, 2H), 7.64 (t, $J$ = 6.8 Hz, 1H), 7.55 (m, 4H), 7.44 (d, $J$ = 7.2 Hz, 2H), 7.22 (m, 2H), 7.11 (m, 2H), 5.89 (s, 1H), 5.00 (s, 1H), 4.53 (d, $J$ = 5.6 Hz, 2H), 3.77 (s, 2H), 3.63 (s, 4H), 3.53 (m, 2H), 3.39 (m, 2H), 3.11 (s, 3H), 1.39 (s, 9H). $^{13}$C NMR (150 MHz, $CDCl_3$): $\delta$ 167.5, 167.4, 167.3, 157.5, 149.1, 141.3, 136.8, 129.7, 129.4, 129.1, 129.0, 128.4, 128.3, 128.2, 127.4, 127.2, 127.0, 123.2, 123.1, 122.4, 122.3, 119.0, 116.4, 58.5, 57.5, 57.1, 46.2, 42.0, 40.0, 28.3, 21.2.

**Zin(1):** To a solution of **21** (1.3 mg, 1.8 µmol, 1.5 eq) in $CH_2Cl_2$ (1.0 mL), TFA (500 µL) was added. The reaction mixture was allowed to stir at r.t. for 30 min. After confirming the consumption of **21** by TLC, the solvent was evaporated and azoetroped with toluene 3 times. To a solution of Alexa Fluor 647 COOH (1 mg, 1.16 µmol, 1.0 eq) in dry DMF (100 µL), TSTU (0.5 mg, 1.66 µmol, 1.5 eq) and $NEt_3$ (10 µL) were added. The reaction mixture was allowed to stir at r.t. under Ar atmosphere for 1 h. After checking the reaction progress by reversed phase TLC, the reaction mixture was added to the deprotected **21** (amine) with additional 300 µL of dry DMF. After stirring at r.t. for 1h, the reaction solution was diluted with $H_2O$ (10 mM TEAA) and purified by RP-HPLC (A/B = 0/100 to 50/50 over 50 min, A: $CH_3CN$, B: $H_2O$ (10 mM TEAA)). After lyophilization, **Zin(1)** (0.5 µmol, 43 % in 2 steps) was obtained as a blue solid. The amount of **Zin(1)** was measured by UV-Vis ($\varepsilon$ = 270000 at 650 nm in $CH_3OH$). HR-ESI MS *m/z* calcd. for $C_{70}H_{77}N_{11}O_{16}S_4$ $[M-3H]^{2-}$ 727.7222, found 727.7222. The purity of **Zin(1)** was checked by HPLC as below ($CH_3CN/H_2O$ (10 mM TEAA) = 0/100 (0 min) → 50/50 (30 min)).

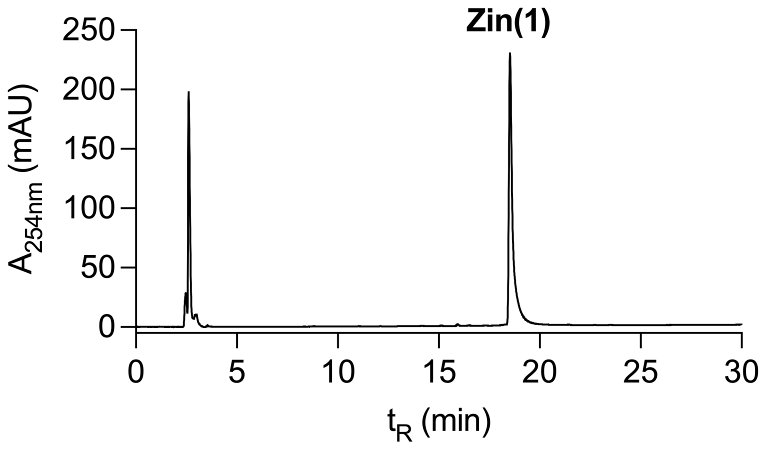
Zin(1)
$A_{254nm}$ (mAU)
250
200
150
100
50
0
0
5
10
15
20
25
30
$t_R$ (min)

#### Synthesis of Zin(2)

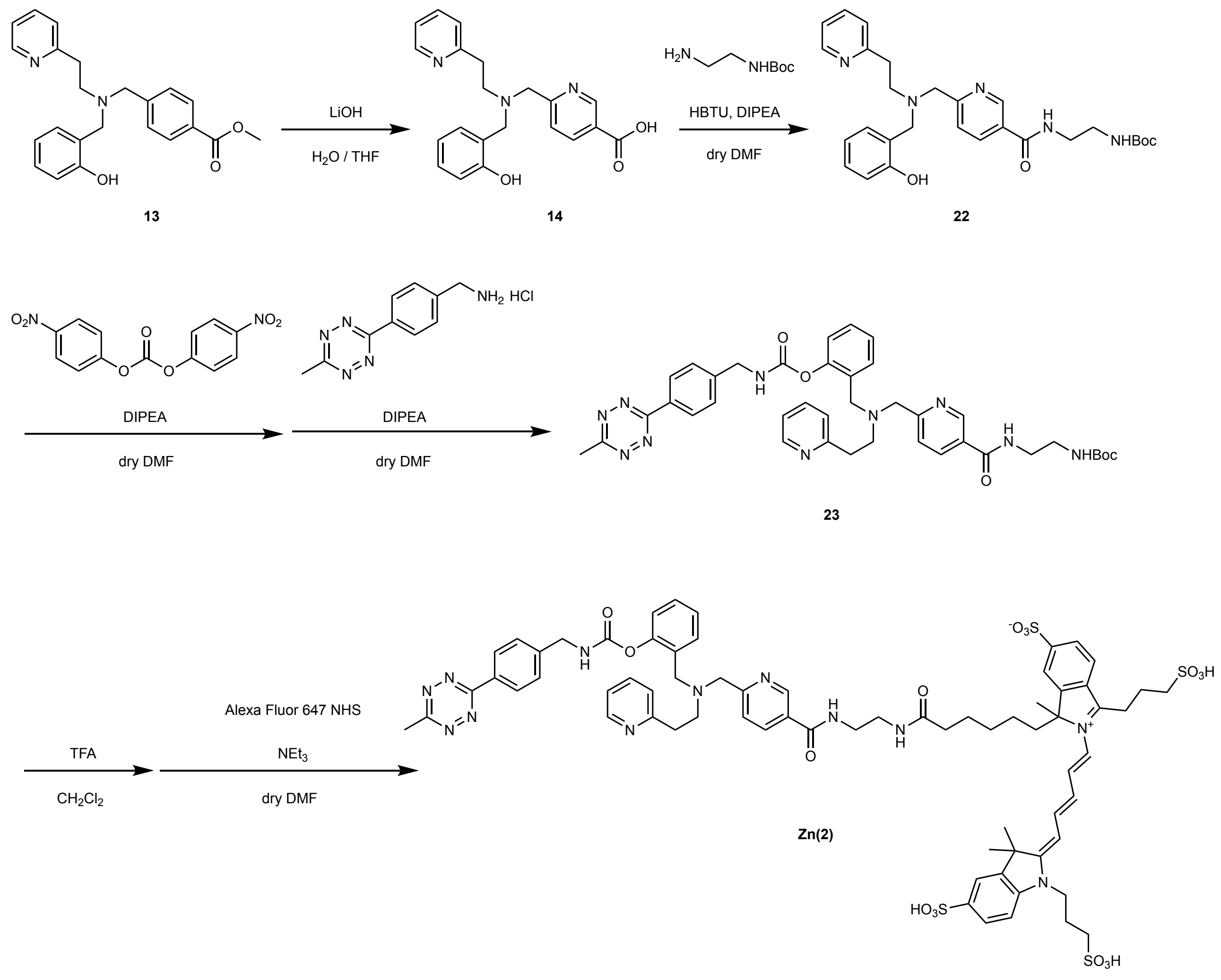


**22:** To a solution of **13** (194 mg, 0.5 mmol, 1.0 eq) in 6 mL of THF/$H_2O$ (1/1, v/v), 1 M LiOH aq. (1.5 mL, 1.5 mmol, 3.0 eq) was added. After stirring at r.t. overnight, the solution was diluted with $H_2O$ and the solution pH was adjusted to 6 with 1 M HCl aq. The solvent was evaporated and the residue was re-dissolved with dry DMF (5 mL). Then, HBTU (300 mg, 0.79 mmol, 1.6 eq), DIPEA (897 µL, 5.1 mmol, 10 eq), and *N*-Boc-ethylenediamine (90.6 mg, 0.6 mmol, 1.2 eq) were added and the reaction mixture was allowed to stir at r.t. overnight. The crude product was purified by flash column chromatography on silica gel (0–17 % $CH_3OH/CHCl_3$) to yield **22** (41.1 mg, 0.081 mmol, 16.2 % in 2 steps) as a yellow oil. $^1H$ NMR (400 MHz, $CDCl_3$): $\delta$ 8.98 (s, 1H), 8.49 (d, $J$ = 4.8 Hz, 1H), 8.04 (dd, $J$ = 8.0, 2.4 Hz, 1H), 7.59 (td, $J$ = 7.6, 1.6 Hz, 2H), 7.20–7.12 (m, 2H), 7.07 (d, $J$ = 7.6 Hz, 1H), 7.03 (d, $J$ = 7.6 Hz, 1H), 6.83 (d, $J$ = 8.0 Hz, 1H), 6.78 (t, $J$ = 7.6 Hz, 1H), 3.89 (s, 2H), 3.86 (s, 2H), 3.60 (m, 2H), 3.44 (m, 2H), 3.05–3.00 (m, 4H), 1.43 (9H, s). $^{13}C$ NMR (150 MHz, $CDCl_3$):

$\delta$ 165.7, 160.6, 159.4, 157.6, 157.4, 149.2, 147.8, 136.6, 135.6, 129.4, 129.0, 128.5, 123.4, 122.9, 122.1, 121.5, 119.2, 116.3, 80.1, 59.1, 57.5, 53.7, 42.2, 40.0, 35.2, 28.3.

**23:** To a solution of **22** (4.6 mg, 9.1 μmol, 1.0 eq) in dry DMF (100 μL), bis(4-nitrophenyl) carbonate (4.1 mg, 13.6 μmol, 1.5 eq) and DIPEA (6.5 μL, 37.2 μmol, 4.0 eq) were added. After stirring at r.t. for 2 h, MeTz-amine HCl salt (3.2 mg, 13.6 μmol, 1.5 eq) and DIPEA (2.6 μL, 15.3 μmol, 1.7 eq) were added. The reaction mixture was allowed to stir at r.t. overnight. After evaporating the solvent, the crude product was purified by flash column chromatography on silica gel (0–10 % $CH_3OH/CHCl_3$) to yield **23** (3.4 mg, 4.6 μmol, 51 %) as a pink solid. $^1$H NMR (400 MHz, $CDCl_3$): $\delta$ 8.92 (1H, s), 8.58 (m, 3H), 8.26 (s, 1H), 8.00 (dd, $J$ = 8.0, 2.0 Hz, 1H), 7.64 (d, $J$ = 8.8 Hz, 2H), 7.54 (d, $J$ = 7.6 Hz, 2H), 7.39 (d, J = 8.4 Hz, 2H), 7.15 (m, 1H), 7.08 (m, 1H), 7.01 (m, 1H), 4.63 (d, $J$ = 6.0 Hz, 2H), 3.85 (s, 2H), 3.78 (s, 2H), 3.58 (m, 2H), 3.43 (m, 2H), 3.12–3.11 (m, 7H), 1.44 (s, 9H). $^{13}$C NMR (150 MHz, $CDCl_3$): $\delta$ 167.3, 163.9, 157.5, 155.3, 149.9, 149.2, 148.3, 147.5, 143.8, 136.6, 135.2, 131.8, 129.4, 129.0, 128.5, 128.4, 128.2, 128.1, 125.2, 123.3, 122.8, 121.4, 119.2, 116.3, 80.3, 60.2, 59.1, 57.6, 45.0, 42.6, 39.9, 35.3, 28.3, 21.2.

**Zin(2):** To a solution of **23** (1.2 mg, 1.64 μmol, 1.4 eq) in $CH_2Cl_2$ (1.0 mL), TFA (100 μL) was added. The reaction mixture was allowed to stir at r.t. for 30 min. After confirming the consumption of **23** by TLC, the solvent was evaporated and azeotroped with toluene 3 times. To a solution of Alexa Fluor 647 COOH (1 mg, 1.16 μmol, 1.0 eq) in dry DMF (100 μL), TSTU (0.5 mg, 1.66 μmol, 1.4 eq) and TEA (10 μL) were added. The reaction mixture was allowed to stir at r.t. under Ar atmosphere for 1 h. After checking the reaction progress by reversed phase TLC, the reaction mixture was added to the deprotected **23** (amine) with additional 300 μL of dry DMF. After stirring at r.t. for 1h, the reaction solution was diluted with $H_2O$ (0.1% TFA) and purified by RP-HPLC (A/B = 10/90 to 50/50 over 40 min, A: $CH_3CN$ (0.1% TFA), B: $H_2O$ (0.1% TFA)). After lyophilization, **Zin(2)** (0.4 μmol, 34.4 % in 2 steps) was obtained as a blue solid. The amount of **Zin(2)** was measured by UV-Vis ($\varepsilon$ = 270000 at 650 nm in $CH_3OH$). HR-ESI MS *m/z* calcd. for $C_{70}H_{78}N_{12}O_{16}S_4$ $[M-3H]^{2-}$ 735.2276, found 735.2283. The purity of **Zin(2)** was checked by HPLC as below ($CH_3CN$ (0.1% TFA)/$H_2O$ (0.1% TFA) = 0/100 (0 min) → 10/90 (5 min) → 50/50 (30 min)).

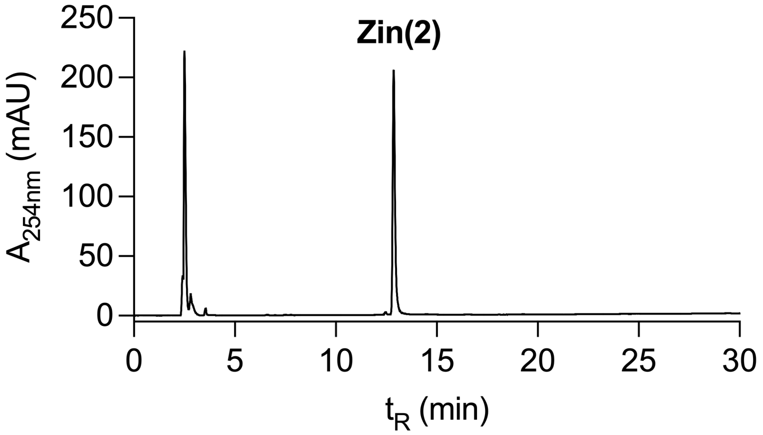
Zin(2)
250
200
150
100
50
0
A254nm (mAU)
0
5
10
15
20
25
30
tR (min)

#### Synthesis of Zin(3)

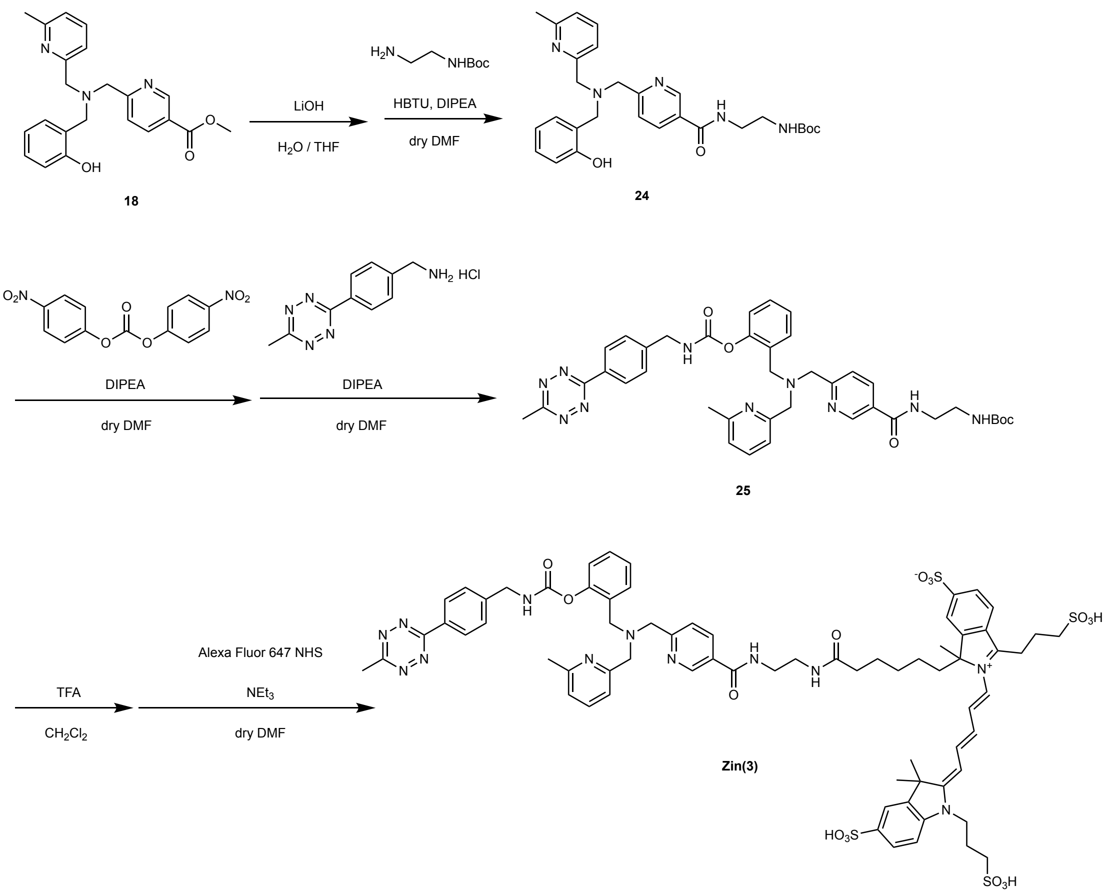


**24:** To a solution of **18** (79 mg, 0.208 mmol, 1.0 eq) in dry $CH_3OH$ (2 mL) and $H_2O$ (1 mL), LiOH•$H_2O$ (34.8 mg, 0.832 mmol, 4.0 eq) was added. After stirring at r.t. for 24 h, the solution was diluted with $H_2O$ and the solution pH was adjusted to 5-6 with 2 M HCl aq. The product was extracted with $CHCl_3$ (10 mL×10). The organic layer was dried over $Na_2SO_4$. After evaporation, the hydrolyzed **18** (71.8 mg, 0.198 mmol, 95.2%) was obtained. The hydrolyzed **18** was re-dissolved with dry DMF (1.5 mL). Then, HBTU (113 mg, 0.297 mmol, 1.5 eq), DIPEA (103 μL, 0.594 mmol, 3.0 eq), and *N*-Boc-ethylenediamine (47.1 μL, 0.297 mmol, 1.5 eq) were added and the reaction mixture was allowed to stir at r.t. for 3 days. The crude product was purified by flash column chromatography on silica gel (0-10 % $CH_3OH/CH_2Cl_2$ and 0-10% $CH_3OH/CHCl_3$ (1% $NH_3$ aq.)) to yield **24** (58.3 mg, 0.115 mmol, 55.3 % in 2 steps) as a colorless oil. $^1$H NMR (400 MHz, $CDCl_3$): $\delta$ 8.97 (s, 1H), 8.06 (dd, $J$ = 8.4, 2.4 Hz, 1H), 7.53–7.47 (m, 3H), 7.16 (t, $J$ = 7.7 Hz, 1H,), 7.08-7.01 (m, 3H), 6.89 (dd, $J$ = 8.2, 1.0 Hz, 1H), 6.77 (td, $J$ = 7.4, 1.2 Hz, 1H), 5.14 (t, $J$ = 5.8 Hz, 1H), 3.89 (s, 2H), 3.83 (s, 2H), 3.79 (s, 2H), 3.56-3.52 (m, 2H), 3.40–3.36 (m, 2H), 2.59 (s, 3H), 1.39 (s, 9H).

**25:** To a solution of **24** (4.7 mg, 9.3 μmol, 1.0 eq) in dry DMF (100 μL), bis(4-nitrophenyl) carbonate (4.2 mg, 13.9 μmol, 1.5 eq) and DIPEA (6.5 μL, 37.2 μmol, 4.0 eq) were added. After stirring at r.t. for 2 h, MeTz-amine HCl salt (3.3 mg, 13.9 μmol, 1.5 eq) and DIPEA (2.5 μL, 13.9 μmol, 1.5 eq) were added. The reaction mixture was allowed to stir at r.t. for 18.5 h. After evaporating the solvent, the crude product was purified by flash column chromatography on silica gel (0–50 % $CH_3OH/CHCl_3$) to yield **25** (4.0 mg, 5.5 μmol, 59 %) as a purple solid. $^1$H NMR (500 MHz, $CD_3OD$): $\delta$ 8.73 (s, 1H), 8.42 (d, $J$ = 8.5 Hz, 2H), 7.99 (d, $J$ = 8.5 Hz, 1H), 7.59 (d, $J$ = 8.5 Hz, 1H), 7.55–7.48 (m, 4H), 7.35 (d, $J$ = 8.0 Hz, 1H), 7.21-7.10 (m, 2H), 7.01 (d, $J$ = 8.0 Hz, 2H), 4.41 (s, 2H), 3.71 (s, 2H), 3.69 (s, 2H), 3.62 (s, 2H), 3.36-3.33 (m, 2H), 3.19–3.16 (m, 2H), 2.95 (s, 3H), 2.38 (s, 3H), 1.30 (s, 9H).

**Zin(3):** To a solution of **25** (1.3 mg, 1.8 μmol, 1.5 eq) in $CH_2Cl_2$ (400 μL), TFA (100 μL) was added. The reaction mixture was allowed to stir at r.t. for 1 h. After confirming the consumption of **25** by TLC, the solvent was evaporated. To a solution of the deprotected **25** (amine) in dry DMSO (100 μL), Alexa Fluor 647 NHS (1.5 mg, 1.2 μmol, 1.0 eq) and DIPEA (30.8 μL, 177 μmol, 150 eq) were added. After stirring at r.t. for 24 h, the reaction solution was diluted with $CH_3CN$ (0.1% TFA)/$H_2O$ (0.1% TFA) and purified by RP-HPLC (A/B = 10/90 to 52/48 over 24 min, A: $CH_3CN$ (0.1% TFA), B: $H_2O$ (0.1% TFA)). After lyophilization, **Zin(3)** (1.0 mg, 0.7 μmol, 58 % in 2 steps) was obtained as a blue solid. HR-ESI MS *m/z* calcd. for $C_{70}H_{78}N_{12}O_{16}S_4$ $[M-3H]^{2-}$ 735.2276, found 735.2277. The purity of **Zin(3)** was checked by HPLC as below. ($CH_3CN/H_2O$ (10 mM TEAA) = 0/100 (0 min) → 50/50 (50 min)).

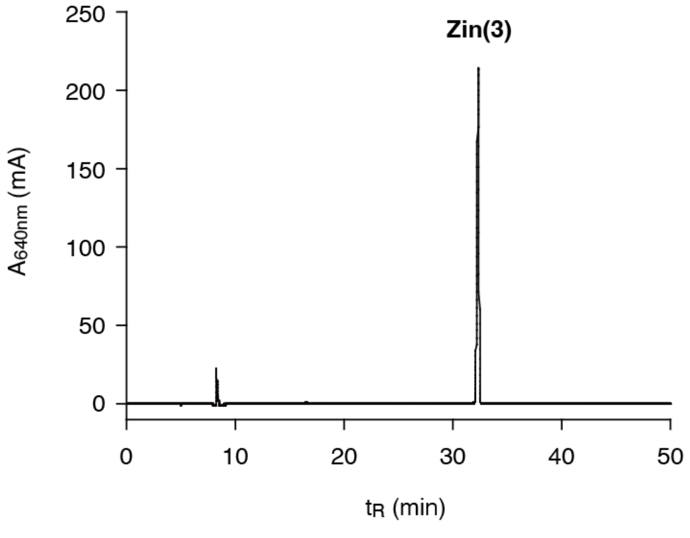

### Synthesis of Fmz (TCO, Ax555)

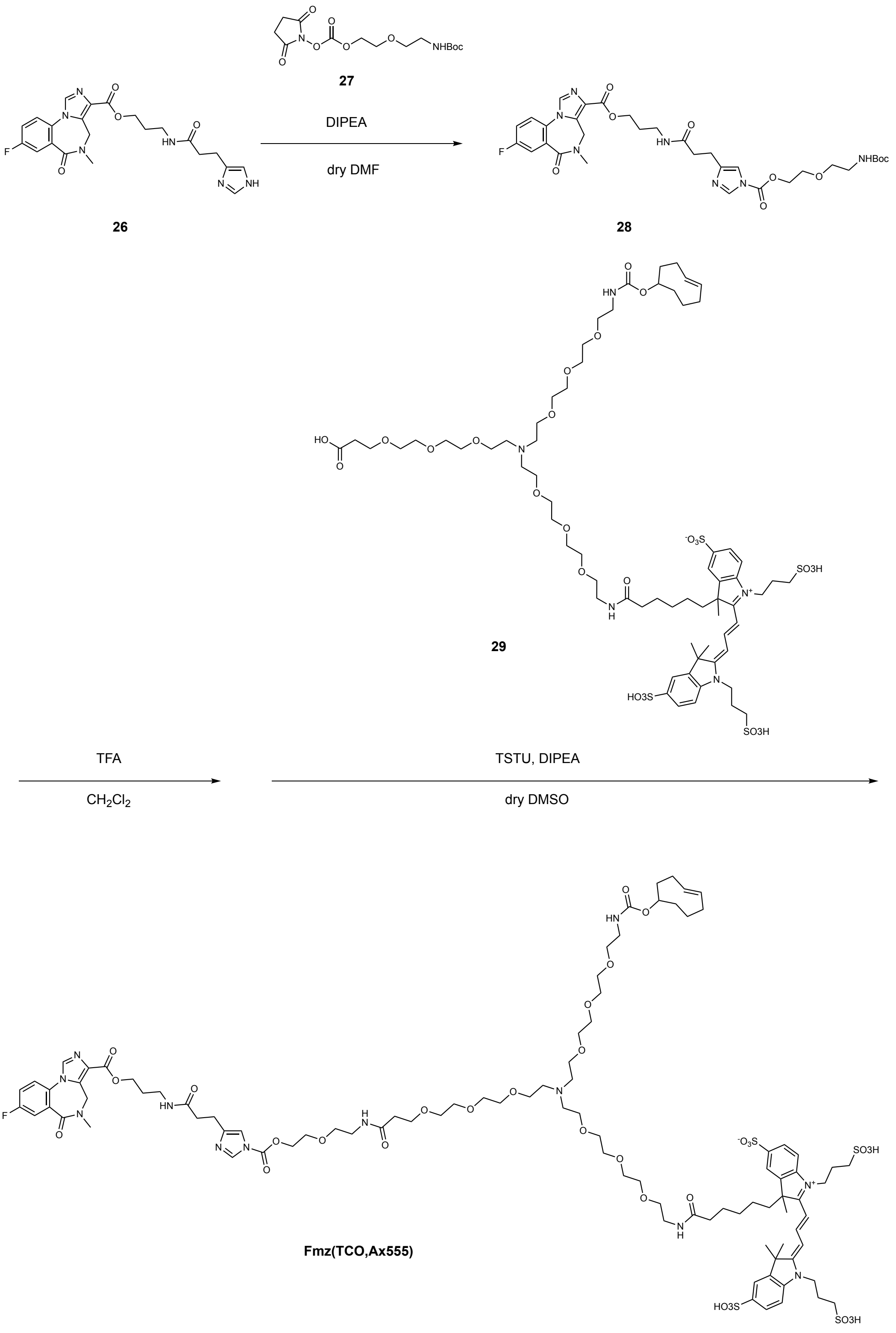

**28**: To a solution of compound **26**[S4] (17.9 mg, 39.4 μmol) in dry DMF (0.5 mL), compound **27**[S5] (27.3 mg, 78.8 μmol, 2.0 eq.) and DIPEA (34 μL, 195 μmol, 5.0 eq.) were added and the reaction mixture was stirred at r.t. under Ar for 3 h. After the evaporation, the residual DMF was azeotropically removed with CH3CN/toluene (1/1) (1 mL×1). The residue was purified with a silica gel chromatography with a linear gradient of MeOH/$CH_2Cl_2$ (5-100%) to give a compound **28** (14.9 mg, 21.7 mmol, 55%). $^1$H NMR(400MHz, $CD_3OD$) δ 8.28 (s, 1H), 8.18 (d, $J$ = 1.2Hz, 1H), 7.75 (dd, $J$= 9.2, 4.8Hz, 1H), 7.70 (dd, $J$= 9.0, 2.9 Hz, 1H), 7.52 (m, 1H) ,7.29 (d, $J$ = 0.8 Hz, 1H), 5.17 (d, $J$ = 14.6 Hz, 1H), 4.50 (m, 3H), 4.31 (bs ,2H), 3.77 (m, 2H), 3.53 (t, $J$ = 5.6 Hz, 2H), 3.35 (t, $J$ = 6.4 Hz, 2H), 3.22 (s, 3H), 3.20 (t, $J$ = 5.6 Hz, 2H), 2.84 (t, $J$ = 7.8 Hz, 2H) ,2.53 (t, $J$ = 7.4 Hz, 2H), 1.95 (m, 2H), 1.39 (s, 9H). HR ESI-MS (calc. for $C_{32}H_{40}FN_7O_9$): $[M + Na]^+$ = 708.2754 (calc. 708.2764).

**Fmz (TCO, Ax555):** To a solution of compound **28** (1.3 mg, 90 μmol) in $CH_2Cl_2$ (0.2 mL), TFA (50 μL) was added. After stirring at r.t. for 1 h, TFA was azeotropically removed with CH3CN / toluene (1/1) (0.4 mL×3). To a solution of compound **29**[S6] (0.4 mg, 0.26 μmol) in dry DMSO (200 μL), TSTU (117 μg, 0.39 μmol, 1.5 eq) and DIPEA (0.5 μL, 2.87 μmol, 11 eq.) were added. The reaction mixture was allowed to stir at r.t. under Ar atmosphere for 1 h. The reaction mixture and additional DIPEA (1.5 μL, 8.62 μL) were added to the Boc-deprotected compound **28** (amine) dissolved in 300 μL of dry DMSO. After stirring at r.t. overnight, the reaction solution was diluted with $H_2O$ (0.1% TFA) and purified by RP-HPLC (($CH_3CN/H_2O$ (10 mM TEAA) = 0/100 (0 min) → 50/50 (50 min)). The compound conferred 2 peaks due to the presence of diastereomers originate from the fluorophore.

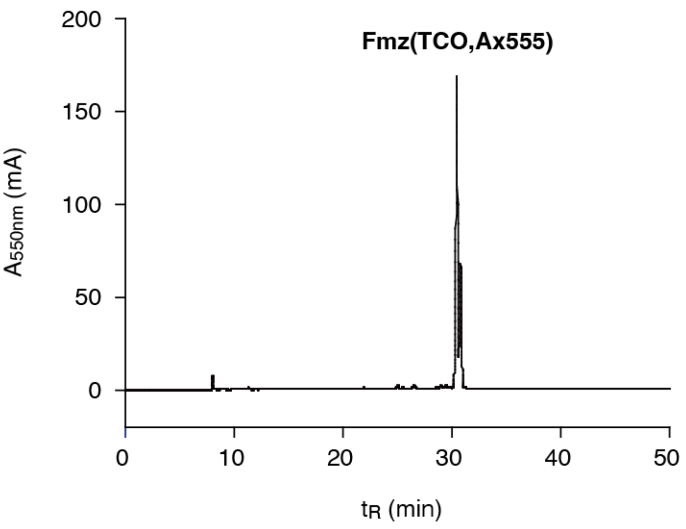
